# Neural Correlates of Optimal Multisensory Decision Making

**DOI:** 10.1101/480178

**Authors:** Han Hou, Qihao Zheng, Yuchen Zhao, Alexandre Pouget, Yong Gu

**Affiliations:** Institute of Neuroscience, Key Laboratory of Primate Neurobiology, CAS Center for Excellence in Brain Science and Intelligence Technology, Chinese Academy of Sciences, Shanghai, China; University of Chinese Academy of Sciences, Beijing, China; University of Geneva, Geneva, Switzerland

**Keywords:** Perceptual decision making, multisensory integration, LIP, probabilistic population code, vestibular, optic flow, self-motion perception

## Abstract

Perceptual decisions are often based on multiple sensory inputs whose reliabilities rapidly vary over time, yet little is known about how our brain integrates these inputs to optimize behavior. Here we show multisensory evidence with time-varying reliability can be accumulated near optimally, in a Bayesian sense, by simply taking time-invariant linear combinations of neural activity across time and modalities, as long as the neural code for the sensory inputs is close to an invariant linear probabilistic population code (ilPPC). Recordings in the lateral intraparietal area (LIP) while macaques optimally performed a vestibular-visual multisensory decision-making task revealed that LIP population activity reflects an integration process consistent with the ilPPC theory. Moreover, LIP accumulates momentary evidence proportional to vestibular acceleration and visual velocity which are encoded in sensory areas with a close approximation to ilPPCs. Together, these results provide a remarkably simple and biologically plausible solution to optimal multisensory decision making.

## Introduction

Most perceptual decisions are based on multiple sensory inputs whose reliabilities vary over time. For instance, a predator can rely on both auditory and visual information to determine when and where to strike a prey, but these two sources of information are not generally equally reliable, nor are their reliabilities constant over time: as the prey gets closer, the quality of the image and sound typically improves, thus increasing their reliabilities. Although such multisensory decision making happens frequently in the real world, the underlying neural mechanisms remain largely unclear.

The so-called drift-diffusion model (DDM) **(^1^Ratcliff, 1978; ^2^Ratcliff and McKoon, 2008; ^3^Ratcliff and Rouder, 1998; ^4^Ratcliff and Smith, 2004)**, a widely used model of perceptual decision making, cannot deal with such decisions optimally in its most standard form. DDMs have been shown to implement the optimal policy for decisions involving just one source of sensory evidence whose reliability is constant over time **(^5^Laming, 1968; ^6^Bogacz, *et al.*, 2006)**. Under such conditions, DDMs can implement the optimal strategy by simply summing evidence over time until an upper or lower bound, corresponding to the two possible choices, is hit **(^6^Bogacz, *et al.*, 2006)**. This type of models lends itself to a straightforward neural implementation in which neurons simply add their sensory inputs until they reach a preset threshold **(^2^Ratcliff and McKoon, 2008; ^7^Gold and Shadlen, 2007)**.

When multiple sensory inputs are involved, the standard DDMs can accumulate sensory evidence optimally as long as the reliabilities of the evidence stay constant during a single trial and across trials. Under this scenario, optimal integration of evidence over time can be achieved by first taking a weighted sum of the momentary evidence at each time step, with weights proportional to the reliability of each sensory stream, followed by temporal integration **(^8^Drugowitsch, *et al.*, 2014)**. However, this strategy no longer works when the reliabilities change over time within a single trial.

In this case, the momentary evidence must be linearly combined with weights proportional to the time-varying reliabilities, which requires that the synaptic weights change on a very fast time scale since, in the real life, reliability can change significantly over tens of milliseconds. Moreover, when the reliabilities of the sensory inputs are not known in advance, which is typically the case in real-world situations, neurons cannot determine how to appropriately modulate their synaptic weights until after the sensory inputs have been observed. Therefore, even if it is possible to extend standard DDMs to time-varying reliability **(^8^Drugowitsch, *et al.*, 2014)**, it is unclear how such a solution could be implemented biologically.

In contrast, there exists another class of models which does not necessarily involve changes in synaptic strength. As long as the sensory inputs are encoded with what is known as “invariant linear probabilistic population codes” (ilPPC), the neural solution for optimal multisensory integration is remarkably simple: it only requires that neurons compute linear combinations of their inputs across time or modalities using fixed—reliability-independent—synaptic weights **(^9^Beck, *et al.*, 2008; ^10^Ma, *et al.*, 2006)**. This solution relies on one specific property of ilPPC: the reliability of the neural code is proportional to the amplitude of the neural responses. As a result, when summing two sensory inputs with unequal reliability, the sensory input with the lowest reliability contribute less to the sum because of its lower firing rate. This is formally equivalent to weighting Gaussian samples with their reliability in an extended DDM, except that there is no need for actual weight changes with ilPPC **(^10^Ma, *et al.*, 2006)**. Hence, the ilPPC framework is a promising solution to multisensory decision-making tasks, but it lacks physiological supports.

To investigate whether the brain may implement this solution, we recorded the activity of single neurons in the lateral intraparietal area (LIP) in macaques trained to discriminate their heading direction of self-motion based on multiple sensory inputs: vestibular signals, visual optic flow, or both. Importantly, the vestibular and visual stimuli followed a Gaussian-shape velocity temporal profile, producing naturally varied cue reliability over time within each trial. This behavioral paradigm has been well-established for studying multisensory heading discrimination in the past decade **(^11^Fetsch, *et al.*, 2012; ^12^Gu, *et al.*, 2008; ^13^Fetsch, *et al.*, 2013)**. Nevertheless, these previous studies focused on areas that encode momentary heading inputs, leaving it unknown how these sensory inputs are further accumulated by downstream neurons (e.g. LIP) during perceptual decision making.

We focus first in LIP because it is the most extensively studied brain region where buildup choice-related activity has been found during visuomotor decisions in macaques **(^7^Gold and Shadlen, 2007; ^14^Shadlen and Newsome, 2001; ^15^Shadlen and Newsome, 1996; ^16^Huk, *et al.*, 2017; ^17^Roitman and Shadlen, 2002)**. In addition, LIP receives abundant anatomical inputs **(^18^Boussaoud, *et al.*, 1990)** from areas encoding momentary vestibular and visual self-motion information for heading discrimination, such as the dorsal medial superior temporal (MSTd) area **(^12^Gu, *et al.*, 2008; ^19^Gu, *et al.*, 2006)** and the ventral intraparietal area (VIP) **(^20^Chen, *et al.*, 2011c; ^21^Chen, *et al.*, 2013)**. It is therefore expected that the activity of LIP neurons should carry buildup choice signals germane to the formation of multisensory decisions. Note that two recent rodent studies **(^22^Nikbakht, *et al.*, 2018; ^23^Raposo, *et al.*, 2014)** also have described multisensory decision signals in rat posterior parietal cortex, a region analogous to its primate counterpart. However, these studies did not characterize the computational solution implemented by these neural circuits, which is precisely the question we investigate here. Specifically, we explored whether the response of LIP neurons is consistent with the ilPPC theory in which neurons take fixed linear combinations of their sensory inputs without any need for complex, time-dependent, modality-specific, reweighting of the sensory inputs during multisensory decision making.

## Results

### Optimal multisensory decision-making behavior on macaques

We trained two macaque monkeys to perform a vestibular-visual multisensory decision-making task **(^12^Gu, *et al.*, 2008)** (**Figure 1a**). On each trial, the monkeys experienced a 1.5s-fixed-duration forward motion with a small deviation either to the left or to the right of the dead ahead. At the end of the trial, the animals were required to report the perceived heading direction by making a saccade decision to one of the two choice targets (**Figure 1b**). We randomly interleaved three cue conditions over trials: a vestibular condition and a visual condition in which heading information was solely provided by inertial cues and optic flow, respectively, and a combined condition consisting of congruent vestibular and visual cues. Importantly, both the vestibular and visual stimuli followed a Gaussian-shape velocity temporal profile, peaking at the middle of the 1.5-s stimulus duration. This modulation of velocity over time has an important implication for the reliability of the sensory inputs provided to the animals. Indeed, previous psychophysical studies have established that a model in which the reliability of the visual flow field is proportional to velocity and the reliability of the vestibular signal is proportional to the acceleration, provides the best fits to the behavioral data **(^8^Drugowitsch, *et al.*, 2014)**. Therefore, this stimulus allows us to test how neural circuits accumulate multisensory evidence whose reliability varies over time with distinct temporal profiles (see below).

**Figure 1.**
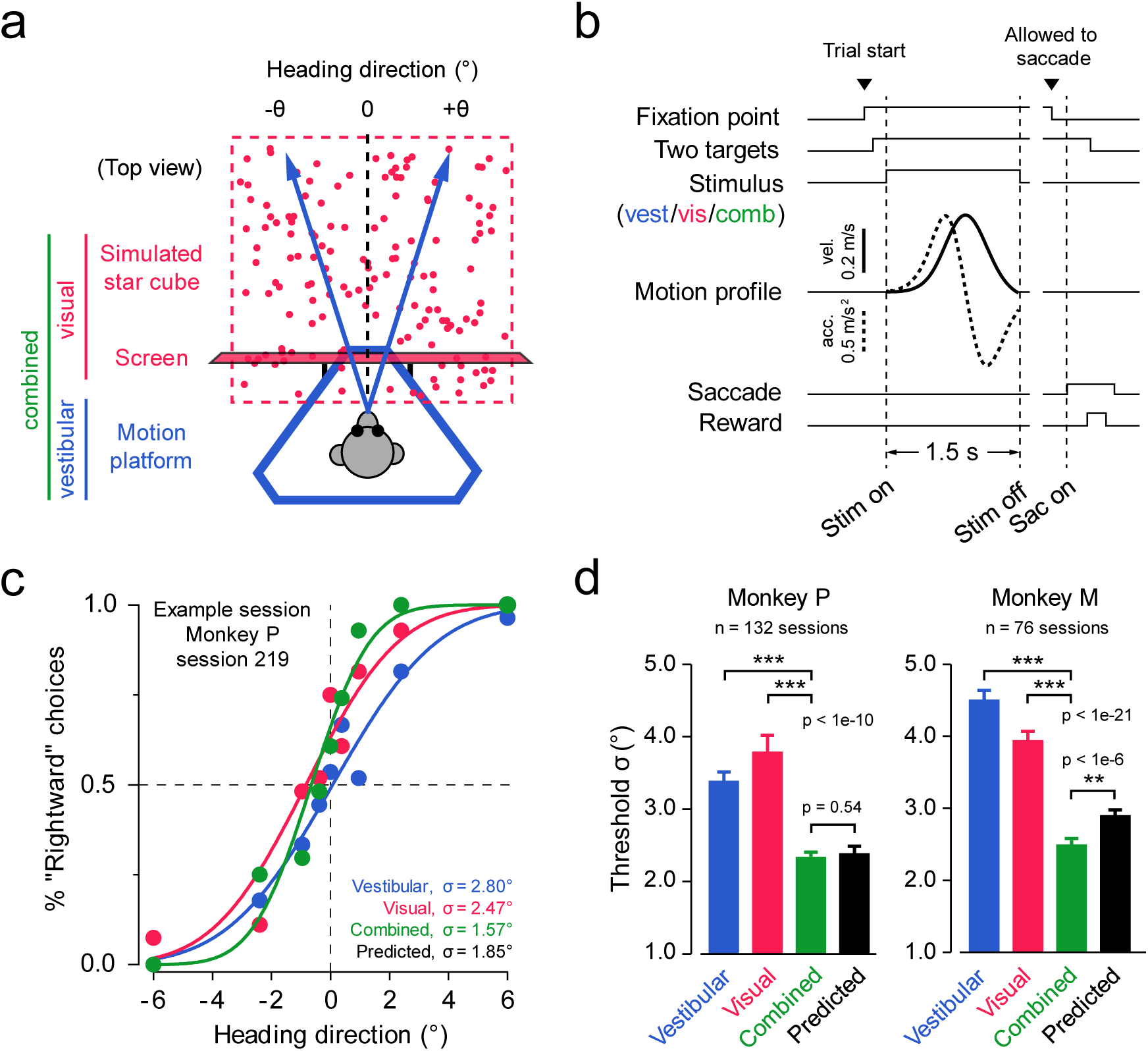
Optimal cue integration in vestibular-visual multisensory decision-making task. **(a)** Schematic drawing of the experimental setup (top view). The vestibular (blue) and visual (red) stimuli of self-motion were provided by a motion platform and an LCD screen mounted on it, respectively. The monkey was seated on the platform and physically translated within the horizontal plane (blue arrows), whereas the screen rendered optical flow simulating what the monkey would see when moving through a three-dimensional star field (red dots). In a combined condition (green), both vestibular and visual stimuli were presented synchronously. The monkey’s task was to discriminate whether the heading direction was to the left or the right of the straight ahead (black dashed line). **(b)** Task timeline. The monkey initiated a trial by fixating at a fixation point, and two choice targets appeared. The monkey then experienced a 1.5-s forward self-motion stimulus with a small leftward or rightward component, after which the monkey reported his perceived heading by making a saccadic eye movement to one of the two targets. The self-motion speed followed a Gaussian-shape profile. **(c)** Example psychometric functions from one session. The proportion of “rightward” choices is plotted against the headings for three cue conditions respectively. Smooth curves represent best-fitting cumulative Gaussian functions. **(d)** Average psychophysical thresholds from two monkeys for three conditions and predicted thresholds calculated from optimal cue integration theory (black bars). Error bars indicate s.e.m.; p values were from paired t-test.

To quantify the monkeys’ behavioral performance, we plotted psychometric curves for each cue condition (**Figure 1c**). Consistent with the previous results **(^12^Gu, *et al.*, 2008)**, the monkeys made more accurate decisions in the combined condition, as evidenced by a steeper psychometric function and a smaller psychophysical threshold (**Figure 1c**). Across all recording sessions and for both monkeys, the psychophysical threshold of the combined condition was significantly smaller than those of single cue conditions and close to the threshold predicted by optimal Bayesian multisensory integration **(^24^Knill and Richards, 1996)** (**Figure 1d**). Therefore, the monkeys can integrate vestibular and visual cues near-optimally during our multisensory decision-making task.

### Heterogeneous multisensory choice signals in LIP

Next, we set out to explore how these optimal decisions were formed in the brain. We recorded from 164 single, well-isolated neurons in LIP of two monkeys while they were performing the task (**Supplementary Figure 1**). As expected, we found buildup choice-related signals in LIP neurons under all cue conditions. As shown in PSTHs of the example cells (**Figure 2a** and **Supplementary Figure 2**), there was generally an increasing divergence between the neuron’s firing rate on trials in which the monkey chose the target in the neuron’s response field (IN choices, solid curves) and trials in which the opposite target was chosen (OUT choices, dashed curves). Importantly, in all cue conditions, the buildup choice signals tended to be stronger for heading directions more distant away from straight ahead (**Supplementary Figure 3**), suggesting that the response of LIP neurons reflects the accumulation of visual and vestibular sensory evidence for heading judgments.

**Figure 2.**
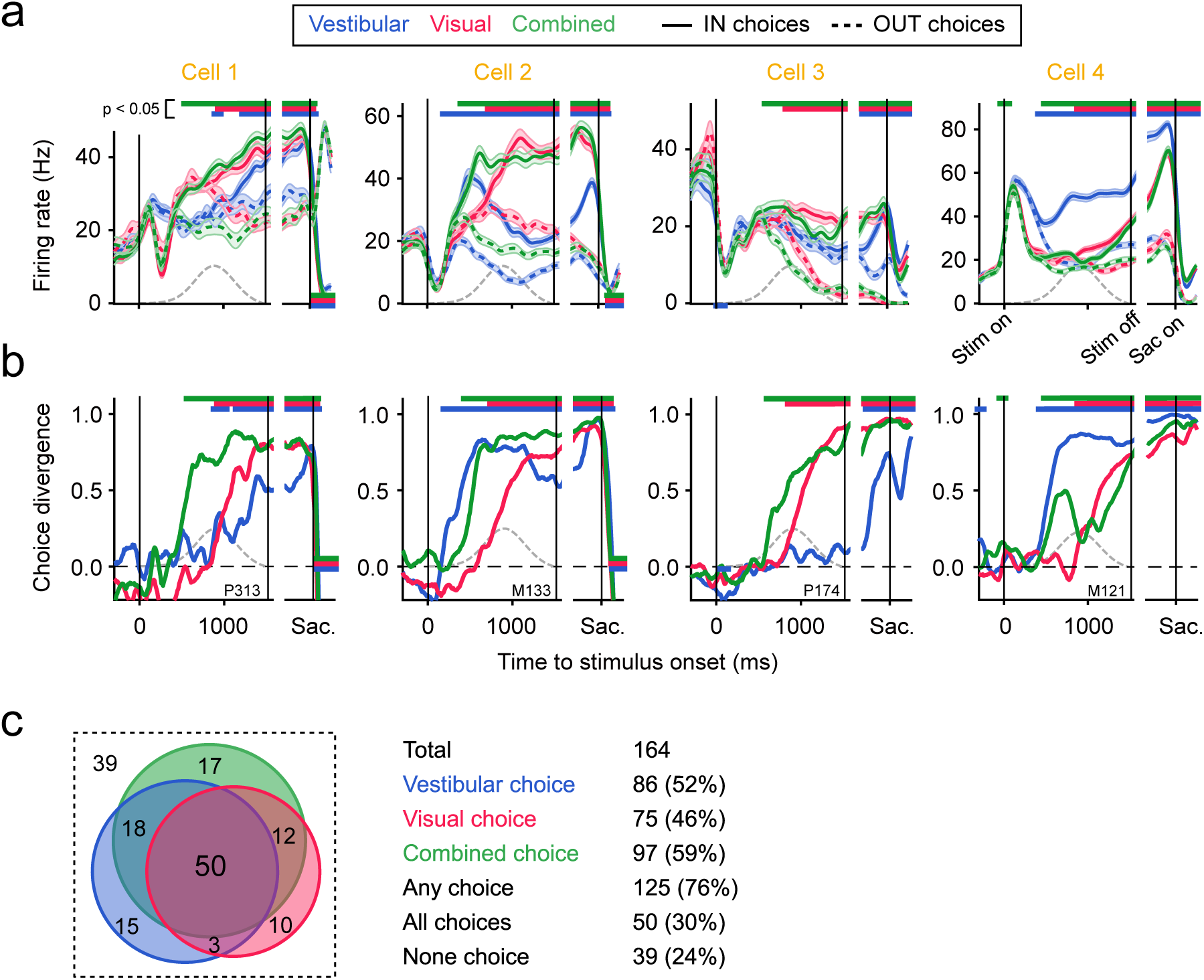
Heterogeneous choice signals in LIP population. **(a)** Peri-stimulus time histograms (PSTHs) of four examples cells. Spike trains were aligned to stimulus onset (left subpanels) and saccade onset (right subpanels), respectively, and grouped by cue condition and monkey’s choice. Vestibular, blue; visual, red; combined, green. Toward the cell’s response field (RF), or IN choices, solid curves; away from the cell’s RF, or OUT choices, dashed curves. Mean firing rates were computed from 10-ms time windows and smoothed with a Gaussian (σ = 50 ms); only correct trials or 0*°* heading trials were included. Shaded error bands, s.e.m. Horizontal color bars represent time epochs in which IN and OUT trials have significantly different firing rates (p < 0.05, t-test), with the color indicating cue condition and the position indicating the relationship between IN and OUT firings (IN > OUT, top; IN < OUT, bottom). Gray dashed curves represent the actual speed profile measured by an accelerometer attached to the motion platform. **(b)** Choice divergence (CD) of the same four cells. CD ranged from −1 to 1 and was derived from ROC analysis for PSTHs in each 10-ms window (see Methods). Horizontal color bars are the same as in **a** except that p-values were from permutation test (n = 1000). **(c)** Venn diagram showing the distribution of choice signals. Numbers within colored areas indicate the numbers of neurons that have significant grand CDs (CD computed from all spikes in 0–1500 ms) under the corresponding combinations of cue conditions.

To better quantify the choice-related signals, we used a ROC analysis to generate an index of choice divergence (CD) **(^23^Raposo, *et al.*, 2014)** that measures the strength of the choice signals (**Figure** 2b). The four cells illustrated in Fig. 2 exhibited canonical ramping choice signals, but their CDs varied greatly across cue conditions. For example, for Cell 1, the CD was largest in the combined condition, modest in the visual condition, and smallest in the vestibular condition. By contrast, for Cell 4, the CD was largest in the vestibular condition. The heterogeneity of choice signals was also manifest at the population level. Approximately half of the LIP neurons exhibited statistically significant CD (p < 0.05, two-sided permutation test) in each cue condition (vestibular: 52%, visual: 46%, combined: 59%; **Figure 2c**), but these three subpopulations did not fully overlap. While more than two thirds of neurons (76%) had significant choice signals in *any* of the three conditions (“Any choice” cells in **Figure 2c**), only a third of neurons (30%) had significant choice signals in *all* of the three conditions (“All choices” cells in **Figure 2c**).

Apart from the heterogeneous choice signals, LIP also encodes heterogeneous sensory modality signals. For example, Cell #12 in **Supplementary Figure 2** exhibited differentiated firing rates across cue conditions without much choice-related signal. In fact, as shown in **Supplementary Figure 4a**, the majority of LIP neurons actually carried mixed choice and modality signals, exhibiting a category-free like neural representation as previously seen in rat posterior parietal cortex **(^23^Raposo, *et al.*, 2014)**. However, although randomly mixed at the single neuron level, the choice and modality signals can still be linearly decoded from the LIP population (**Supplementary Figure 5**). Therefore, we ignore the mixed modality signals thereafter, since they are irrelevant to our heading discrimination task and orthogonal to the decision signals that we really care about.

Another potential difficulty in interpreting LIP activity arises from the fact that LIP neurons also multiplex a combination of temporally overlapping decision-and non-decision-signals **(^25^Park, *et* al., 2014; ^26^Meister, *et al.*, 2013)**. In particular, the signal of saccade preparation may interfere with the one reflecting evidence accumulation **(^14^Shadlen and Newsome, 2001)**. However, this was not likely to be an issue in our study. In our fixed-duration task, we introduced a 300–600 ms delay between the stimulus offset and the time at which the monkey was allowed to saccade (see Methods). Moreover, the monkeys tended to stop integrating evidence around 500 ms prior to the stimulus offset (see **Figure 3b** and below), further separating in time the processes of evidence accumulation and saccade preparation. Therefore, the premotor activity of LIP should not play a significant role in our analysis of multisensory evidence accumulation.

**Figure 3.**
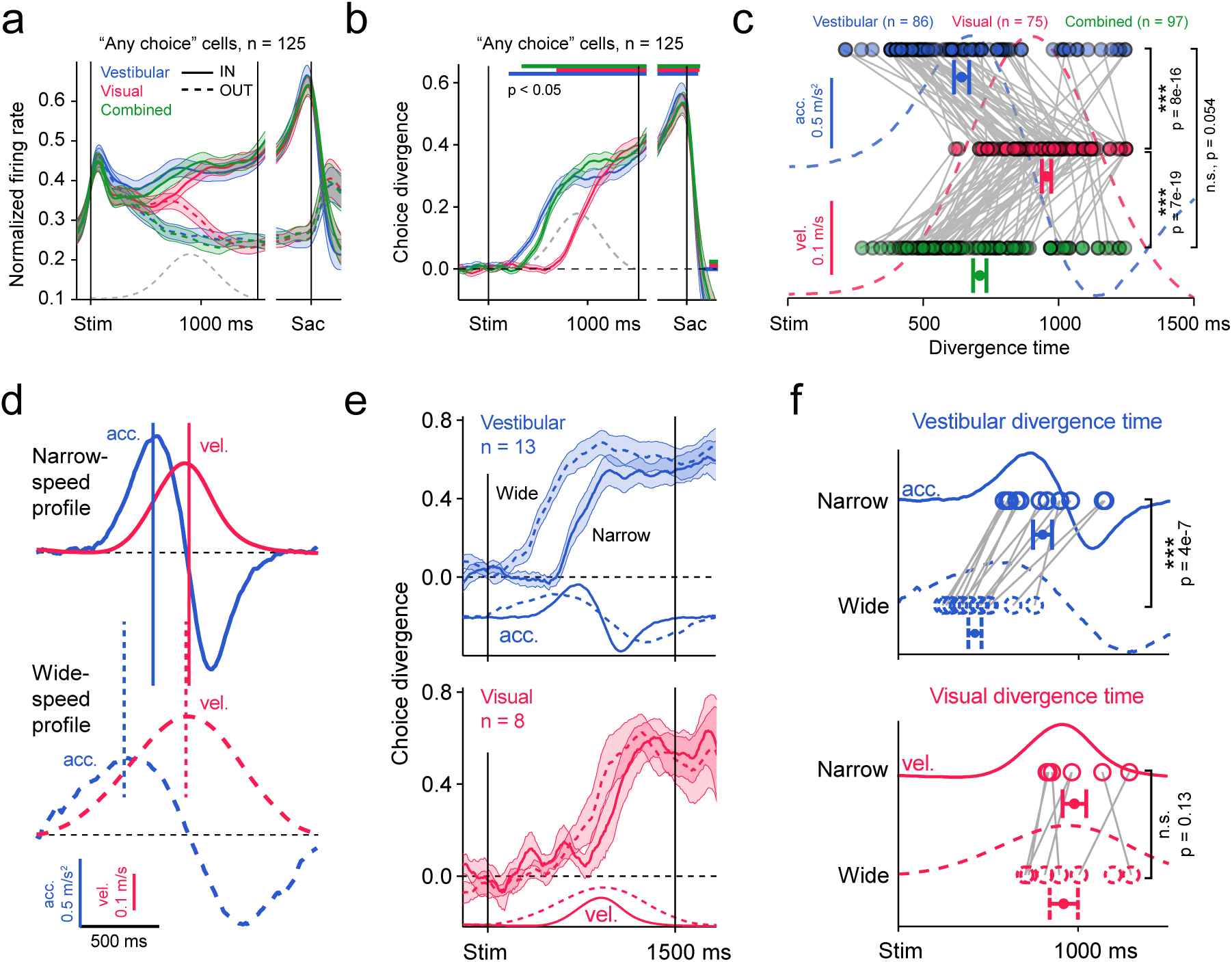
LIP integrates vestibular acceleration but visual speed. **(a and b)** Population average of normalized PSTHs (**a**) and CD (**b**) from 125 “any choice” cells. The vestibular (blue) and combined (green) CDs ramped up much earlier than the visual one (red). Horizontal color bars indicate the time epochs in which population CDs are significantly larger than zero (p < 0.05, t-test). Gray dashed curve, the actual Gaussian speed profile; shaded error bands, s.e.m. **(c)** Divergence time of cells with significant grand CD for each condition. Divergence time was defined as the first occurrence of a 250-ms window in which CD was consistently larger than zero (p < 0.05, permutation test). Gray lines connect data from the same cells; acceleration and speed profiles shown in the background. Data points with horizontal error bars, mean ± s.e.m. of population divergence time; p values, t-test. **(d)** Two motion profiles used to isolate contributions of acceleration and speed to LIP ramping. Top and solid, the narrow-speed profile; bottom and dashed, the wide-speed profile; blue, acceleration; red, speed. Note that by widening the speed profile, we shifted the time of acceleration peak forward (blue vertical lines) while keeping the speed peak unchanged (red vertical lines). **(e)** Vestibular and visual CDs under the two motion profiles. **(f)** Comparison of divergence time between narrow and wide profiles. Note that the vestibular divergence time was significantly shifted, whereas the visual one was not, indicating that LIP integrates sensory evidence from vestibular acceleration and visual speed.

### LIP integrates vestibular acceleration and visual velocity

Despite the high degree of heterogeneity, there was nonetheless a property shared amongst the LIP neurons, namely, the temporal dynamics of the ramping activity was significantly faster in the vestibular and combined conditions than in the visual condition (**Figure 3**). This was evident not only in the averaged rate-based or ROC-based measures (“Any choice” cells, **Figure 3a, b**), but also in the cell-by-cell analysis (**Figure 3c**). Notably, the averaged divergence time under the vestibular and combined conditions aligned well to the acceleration peak of the Gaussian-shape motion profile, whereas the divergence time under the visual condition better aligned to the velocity peak (**Figure 3c, d**ashed curves). This suggests that the physical quantities being integrated over time are speed for the visual stimulus and acceleration for the vestibular stimulus.

An alternative explanation, however, might be that the apparent ∼400 ms interval between the vestibular and visual ramping was caused purely by a difference in their sensory latencies rather than in their underlying physical quantities. For example, LIP activity could have been driven by an ultrafast vestibular signal but a slow visual signal, both of which followed the velocity of the motion. To test this, we designed an experiment in which we used two distinct velocity profiles, a wide one and a narrow one (**Figure 3d**). These profiles were designed to have temporally aligned velocity peaks but misaligned acceleration peaks. If our original physical-quantity hypothesis was correct, we would expect the visual ramping to remain nearly the same under both profiles, while the vestibular ramping should start earlier for the wide profile than for the narrow one, thus reflecting the earlier acceleration peak under the wide profile. In contrast, if the sensory-latency hypothesis was correct, there should be no shift in either the vestibular or visual ramping across the two profiles. Our data matches the first prediction (**Figure 3e, f**). In other words, the temporal discrepancy between the vestibular and visual ramping activities indeed resulted from different physical quantities underlying the momentary evidence fed into LIP. This physiological finding echoed a recent psychophysical study showing that, at the behavioral level, human subjects optimally integrate vestibular and visual momentary evidence with reliability following the amplitude of acceleration and velocity, respectively **(^8^Drugowitsch, *et al.*, 2014)**.

### Network model implementing ilPPC for multisensory decision making

Next, we developed a neural model of multisensory decision making (refer to as M1 thereafter) which takes as input vestibular neurons tuned to acceleration and visual neurons tuned to velocity as observed *in vivo* (equation (2) and (3) in **Methods**; **Figure 4a**). These inputs converge onto an integration layer which takes the sum of the visual and vestibular inputs, as well as integrates this summed input over time. This layer projects in turn to an output layer, labeled LIP, which sums the integrated visuo-vestibular inputs with the activity from another input layer encoding the two possible targets to which the animal can eventually saccade (**Figure 4b, c**). As long as the input layers encode the sensory inputs with ilPPC, this simple network can be shown analytically to implement the Bayes optimal solution even when the reliability of the sensory inputs vary over time as is the case in our experiment **(^9^Beck, *et al.*, 2008; ^10^Ma, *et al.*, 2006)**. Note that separating the integration layer from the LIP layer is not critical to our results. We did so to reflect the fact that current experimental data suggest that LIP may not be the layer performing the integration *per se*, but may only reflect the results of this integration **(^27^Katz, *et al.*, 2016)**.

**Figure 4.**
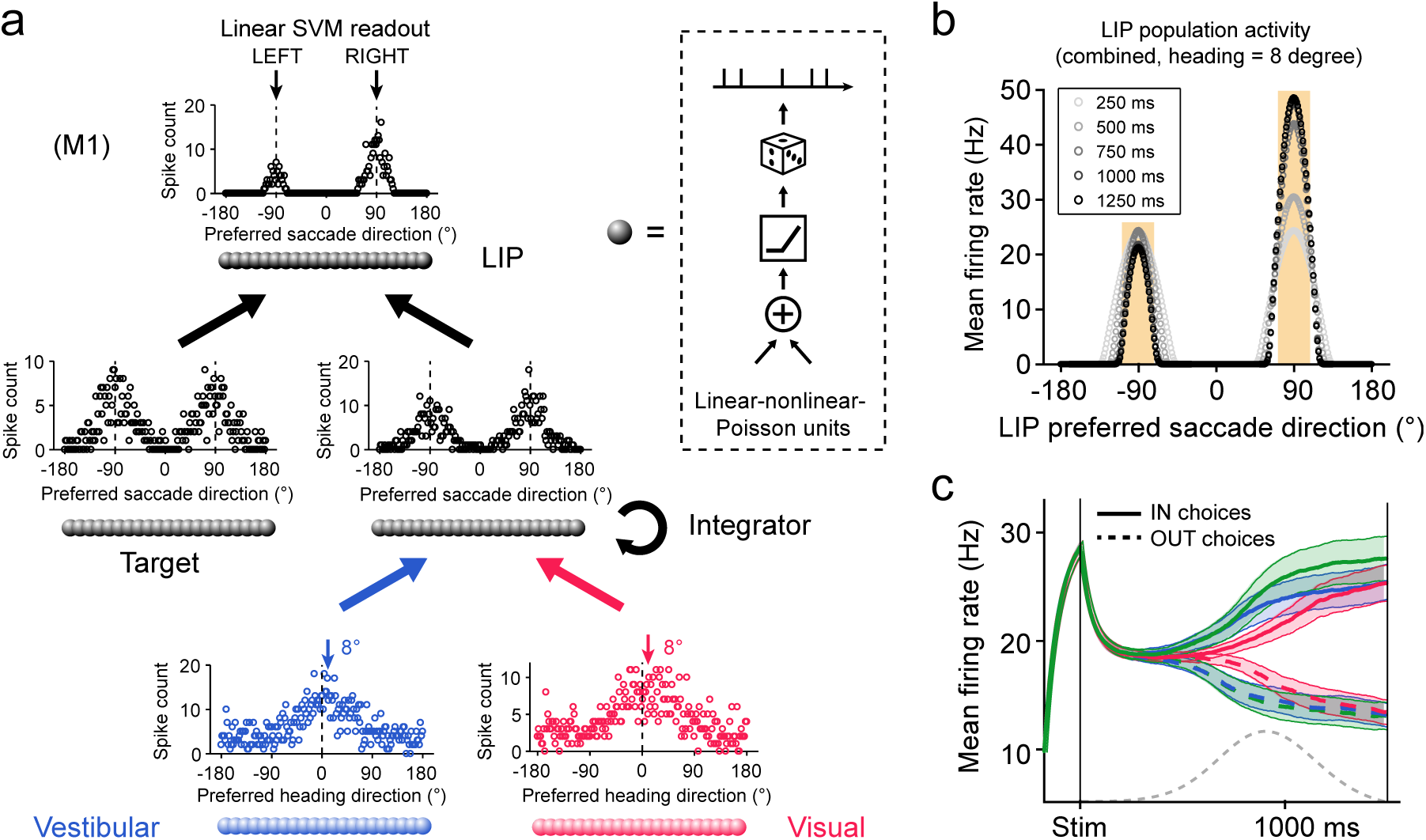
Neural network model with invariant linear probabilistic population codes (ilPPC). **(a)** Network architecture of model M1. The model consists of three interconnected layers of linear-nonlinear-Poisson units (inset). Units in Vestibular and Visual layers have bell-shape ilPPC-compatible tuning curves for heading direction and receive heading stimuli with temporal dynamics following acceleration and speed, respectively. The intermediate Integrator layer simply sums the incoming spikes from the two sensory layers over time and transforms the tuning curves for heading direction to that for saccade direction (−90°, leftward choice; +90°, rightward choice). The LIP layer receives the integrated heading inputs from the Integrator layer, together with visual responses triggered by the two saccade targets. LIP units also have lateral connections implementing short-range excitation and long-range inhibition. Once a decision boundary is hit, or when the end of the trial is reached (1.5 s), LIP activity is decoded by a linear support vector machine for action selection (see **Methods**). Circles indicate representative patterns of activity for each layer; spike counts from 800–1000 ms; combined condition, 8° heading. **(b)** Population firing rate in the LIP layer at five different time points (the same stimulus as in **a**, averaged over 100 repetitions). **(c)** Average PSTHs across LIP population. Trials included three cue conditions and nine heading directions (±8°, ±4°, ±2°, ±1°, 0°). To mimic the experimental procedure, only units with preferred saccade direction close to ±90° were used (with deviation less than 20°; yellow shaded area in **b**). Notations are the same as in **Figure 2a and Figure 3a**.

In an ilPPC, the gain, or amplitude, of the tuning curves of the neurons should be proportional to the reliability of the encoded variable. For instance, in the case of vestibular neurons, the amplitude of the tuning curves to heading should scale with acceleration. In vivo, however, the responses of vestibular and visual neurons are not fully consistent with the assumption of ilPPC because while the amplitude does increase with reliability, in some neurons, the baseline activity decreases with reliability (equation (2) and (3) in **Methods** and **Supplementary Figure 6a**). This violation of the ilPPC assumption implies that a simple sum of activity could incur an information loss. Fortunately, this information loss is small for a population of neurons with tuning properties similar to what has been reported experimentally and information limiting correlations **(^28^Moreno-Bote, *et al.*, 2014)**. Indeed, we found numerically that the information loss was around 5% over a wide range of parameters values (Fano factor, mean correlation, baseline changes, and so on) (**Supplementary Modeling** and **Supplementary Figure 6**).

Importantly, we also endowed the network with a stopping mechanism which terminates sensory integration whenever a function of the LIP population activity reaches a preset threshold (see Methods). Our experiment is not a reaction time experiment and may not require, in principle, such a stopping bound. However, as can be seen in **Figure 3b, L**IP population response saturates around 1s, suggesting that evidence integration stops prematurely. This is indeed consistent with the previous results suggesting that animals and humans use a stopping bound even in fixed duration experiments **(^29^Kiani, *et al.*, 2008)**.

### LIP data are compatible with the ilPPC framework

In the first set of simulations on M1, we adjusted the height of the stopping bounds and found that the model can replicate the near optimal animals’ performance (**Figure 5a**). We then plotted the activity of a typical output neuron (in the LIP layer) in all three conditions. As expected, the activity in the combined condition is roughly equal to the sum of the vestibular-only activity and visual-only activities (**Figure 5b**), at least in the first half of the trial. In the second half of the trial, the activity in the combined condition deviates strongly from the sum because the traces correspond to averages across trials that terminated at different times on different trials due to the stopping bound.

**Figure 5.**
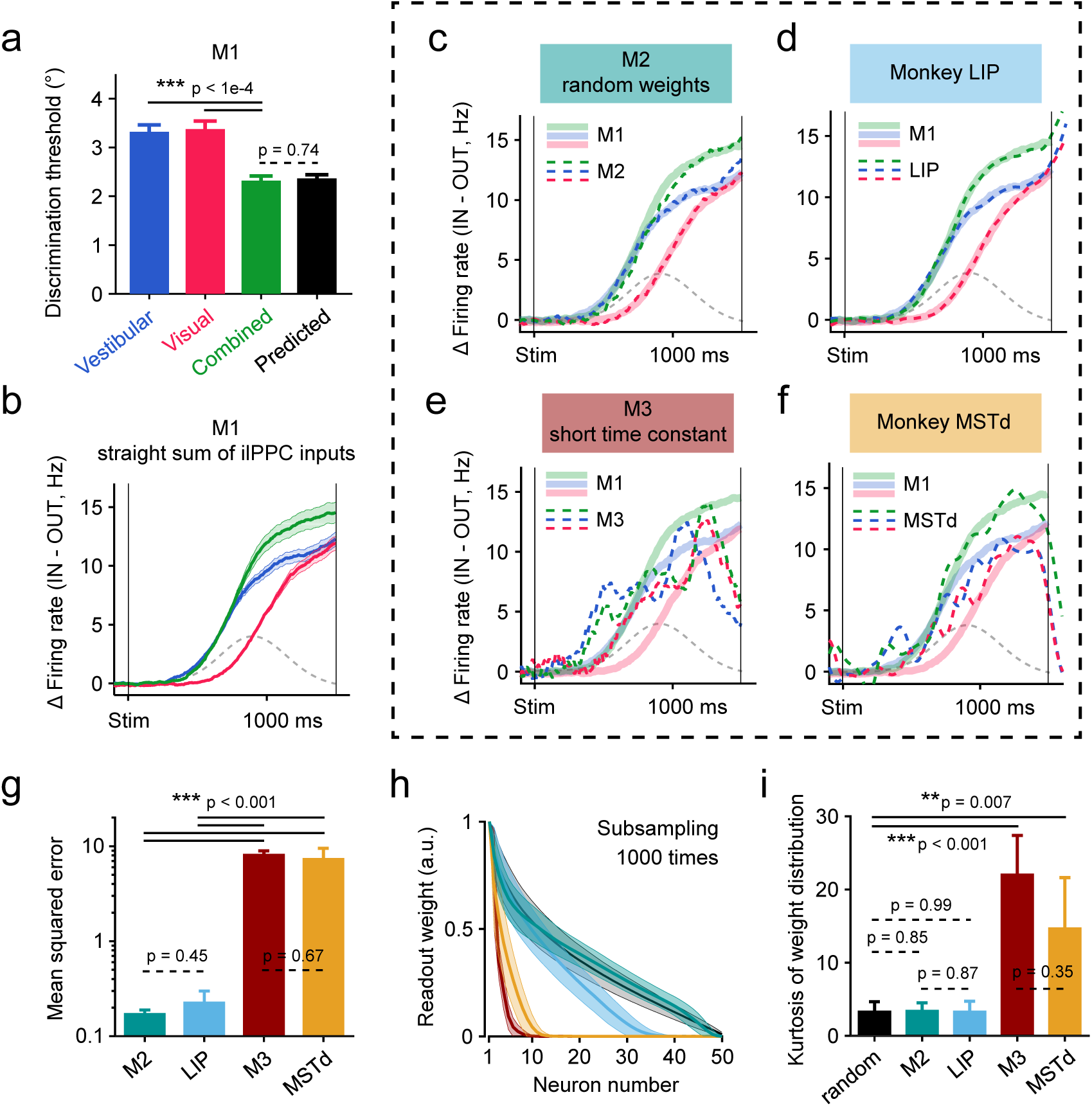
Optimal ilPPC model M1 can be linearly approximated by M2 and LIP but not by M3 and MSTd. **(a)** M1 exhibited near-optimal behavior as the monkey. The psychophysical threshold under the combined condition as indistinguishable from the Bayesian optimal prediction (black). **(b)** Ramping activity of M1 computed as the of PSTHs for IN and OUT trials. Activities from hypothetical units in the LIP layer with preferred direction close ere averaged together (see **Figure 4c** and **Methods**). Since M1 is optimal and homogeneous, we refer to M1’s as “optimal traces” (see the main text). Notations are the same as before. **(c)** Optimal traces from M1 (thick ands) can be linearly reconstructed by population activities obtained from a heterogenous model M2 (dashed Model M2 had the same network architecture as M1 except that it relies on random combinations of ilPPC inputs gration layer (see **Methods**). **(d)** Optimal traces can also be linearly reconstructed by heterogenous single neuron from the LIP data. The similarity between **c** and **d** suggests that both model M2 and monkey LIP are eous variations of to the optimal ilPPC model M1. **(e and f)** In contrast, the optimal traces cannot be cted from activities of a suboptimal model M3 (**e**) or from the MSTd data (**f**), presumably because the time in M3 and MSTd were too short. (**g**) Mean squared error of the fits in panels **c–f**. Error bars and p values were ampling test (n = 50 neurons, 1000 times). (**h**) Normalized readout weights ordered by magnitude. Shaded error icate standard deviations of the subsampling distributions. (**i**) The kurtosis of the distributions of weights. The black curve in (**h**) and black bar in (**e**) were from random readout weights (see **Methods**).

Neurons in M1 are homogeneous in the sense that they all take a perfect sum of their vestibular and visual inputs. Importantly, however, optimal integration does not require such a perfect sum; it can also be achieved with random linear combinations of vestibular and visual inputs **(^10^Ma, *et al.*, 2006)**. Accordingly, we simulated a second model, refer to as M2, in which the visual and vestibular weights of each neuron were drawn from lognormal distributions (**Figure 5c** and see **Methods**). Like M1, model M2 can be tuned to reproduce the Bayes optimal discrimination thresholds (**Supplementary Figure 7a, b**). However, in contrast to model M1, the neurons showed a wide range of response profiles closer to what we observed in vivo (**Supplementary Figure 7c**). In particular, we found that the distribution of visual and vestibular weights was similar in the model and in LIP data (**Supplementary Figure 7d**).

Since model M2 is a linear combination away from model M1, we tested whether the response of M1 neurons could be estimated by linearly combining the response of M2 neurons. Multivariate linear regression confirmed that M1 response profiles could indeed be perfectly reproduced by linearly combining M2 responses (**Figure 5c**). Since LIP neurons also appear to be computing random linear combinations of visual and vestibular inputs, the same result should hold for LIP responses. This is indeed what we found: the response of M1 neurons can be closely approximated by linearly combining the response of LIP neurons (**Figure 5d, g** and Supplementary Figure 9).

This last result is key: it suggests that LIP neurons behave quite similarly to the neurons in M2. The two sets of neurons, however, differ quite significantly in how they integrate their inputs over time. LIP neurons display a wide variety of temporal profiles (see **Supplementary Figure 2**), suggesting that very few neurons act like perfect temporal integrators, in contrast to M2 neurons. Nonetheless, the fact that linear combinations of LIP neurons could reproduce the response of M1 neurons indicates that LIP responses provide a basis set sufficiently varied to allow perfect integration at the population level, a result consistent with what has been recently reported in the posterior parietal cortex of rats engaged in a perceptual decision making task **(^30^Scott, *et al.*, 2017)**.

In addition to this second model, we simulated a third model (M3) in which the time constant of the integration layer was reduced to 100 ms. Interestingly, we found that it was not possible to linearly combine the responses of M3 output neurons to reproduce the traces of the optimal model M1 (**Figure 5e, g**), thus emphasizing the importance of long integration time constant for fitting the optimal model. We also wondered whether M1 could be fitted by the response of MSTd neurons, which are known to combine visual and vestibular responses and whose time constant are believed to be of the same order as model M3. We found that the fit to M1 from MSTd neurons was markedly worse than those obtained from M2 and LIP but was close to that from M3 (**Figure 5f, g**). Moreover, only a small fraction of cells contributed significantly to this fit, in sharp contrast to what we observed in M2 and LIP (**Figure 5h, i**). In fact, the late phase of M1 responses was captured mostly by MSTd cells with short time constants who seemed sensitive to deceleration, rather than integrating cells (**Supplementary Figure 8**).

Finally, we computed the shuffled Fisher information over time for the models and the experimental data (**Figure 6**). The Fisher information in a neuronal population is a measure inversely proportional to the square of the discrimination threshold of an ideal observer **(^31^Beck, *et al.*, 2011; ^32^Seung and Sompolinsky, 1993)**. The shuffled Fisher information is a related measure corresponding to the information in a data set in which neurons are recorded one at a time, as opposed to simultaneously, which is the case for our data set **(^33^Series, *et al.*, 2004)** (see **Methods**). Our network simulations revealed that the shuffled Fisher information should increase over time in all conditions, reflecting the temporal accumulation of evidence (**Figure 6a**). In addition, we observed that this rise in information starts earlier in the vestibular condition than in the visual one because of the temporal offset between acceleration and velocity. In the combined condition, the Fisher information follows at first the vestibular condition before exceeding the vestibular trace once the visual information becomes available. Remarkably, the shuffled Fisher information estimated from the LIP responses follows qualitatively the same trend as the ones observed in the model (**Figure 6b**). In contrast to M2 and LIP neurons, shuffled Fisher information in M3 and MSTd followed the profile expected for neurons with short time constant: it simply reflected the velocity profile of the stimulus and did not exhibit the plateau expected from a decision area (**Figure 6c, d**).

**Figure 6.**
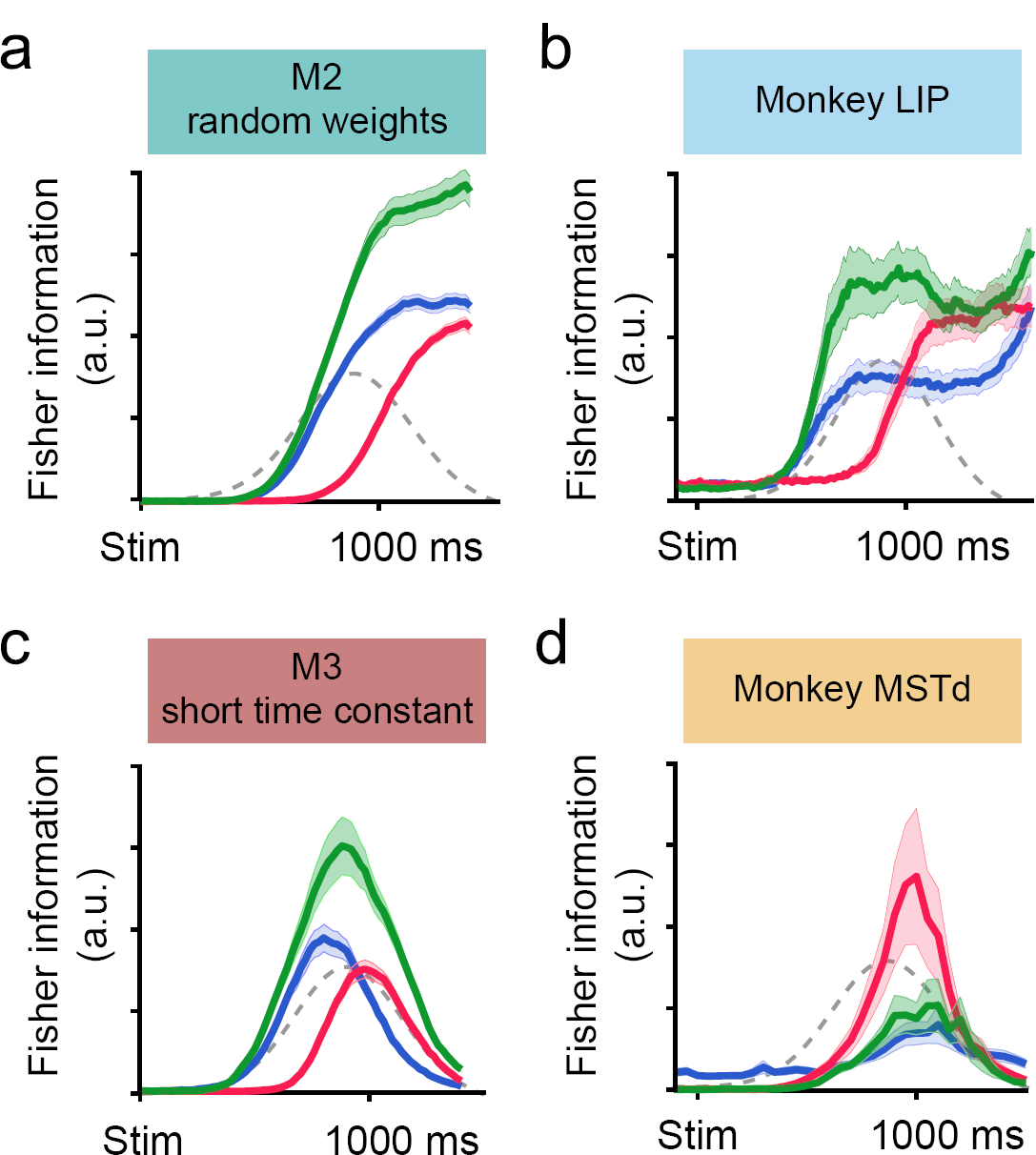
Shuffled Fisher information for the model and the experimental data. **(a)** Shuffled Fisher information of M2 calculated by 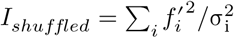, where 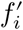 denotes the derivative of the local tuning curve of the th neuron and denotes the averaged variance of its responses around 0° (see **Methods**). Both correct and wrong trials were included. Shaded error bands, s.e.m. estimated from bootstrap. Note that the absolute value of shuffled Fisher information is arbitrary. **(b-d)** Same as in **a** but for the monkey LIP data, the M3 responses, and the monkey MSTd data, respectively. Note that LIP is similar to M2, and MSTd to M3.

Taken together, our results are consistent with the notion that MSTd neurons provide the visual momentary evidence for decision making, while LIP circuits, or circuits upstream from LIP, implement the near optimal solution of model M1, in the sense that the LIP population activity is a mere linear transformation away from that solution.

## Discussion

Integrating ever-changing sensory inputs from different sources across time is crucial for animals to optimize their decisions in a complex environment, yet little is known about the underlying mechanisms, either experimentally or theoretically. In the current study, we present, to the best of our knowledge, the first electrophysiological data on multisensory decision making from non-human primates. We found that LIP neurons in the macaque posterior parietal cortex encode ramping decision signals not only for the visual condition, as widely shown in the literature, but also for the vestibular and combined conditions, except with distinct temporal dynamics. Importantly, these data are compatible with an ilPPC framework where optimal multisensory evidence accumulation is achieved by simply summing sensory inputs across both modalities and time, even with mismatched temporal profiles of cue reliabilities and with heterogeneous sensory-motor representation. Therefore, our results provide the first neural correlate of optimal multisensory decision making.

### Distinct visual and vestibular temporal dynamics in LIP

By comparing the temporal dynamics of LIP population under different modalities, we found that LIP neurons accumulate vestibular acceleration and visual speed, which serve as momentary evidence for their respective modalities. These findings may seem confusing at first glance, since it is more intuitive to assume that neural circuits would combine evidence with the same temporal dynamics across cues, namely, either visual and vestibular speed or visual and vestibular acceleration **(^19^Gu, *et al.*, 2006; ^34^Chen, *et al.*, 2011a; ^35^Fetsch, *et al.*, 2010; ^36^Smith, *et al.*, 2017)**. In support of this idea, recent studies have found a remarkable transformation from acceleration-dominated to speed-dominated vestibular signal along the vestibular pathway, i.e. from peripheral otolith organs to the central nervous system **(^37^Laurens, *et al.*, 2017)**, as well as a moderate but noticeable further transformation along several sensory cortices **(^19^Gu, *et al.*, 2006; ^34^Chen, *et al.*, 2011a**; ^35^Fetsch, *et al.*, 2010; ^37^Laurens, *et al.*, 2017). Given that visual motion responses are typically dominated by speed **(^19^Gu, *et al.*, 2006; ^38^Lisberger and Movshon, 1999)**, one would think that the brain may deliberately turn the vestibular signal from acceleration-to speed-sensitive to facilitate the combination with the visual signal.

However, if the vestibular momentary evidence is proportional to acceleration corrupted by white noise across time, integrating this evidence to obtain a velocity signal would not simplify decision making. On the contrary, this step would introduce temporal correlations **(^39^Churchland, *et al.*, 2011)**, in which case, even with ilPPC, a simple sum of the momentary evidence would no longer be optimal **(^6^Bogacz, *et al.*, 2006)**. Instead, downstream circuits would have to compute a weighted sum of the sensory evidence, which would effectively differentiate the momentary evidence before summing them. In other words, optimal integration would effectively recover the original acceleration signals. Our results, along with previous psychophysical results **(^8^Drugowitsch, *et al.*, 2014)**, strongly suggest that the brain does not go through this extra step and uses the acceleration signals as momentary evidence instead.

### Multisensory convergence in the brain for heading decision

One of the long-standing questions about multisensory integration is whether integration takes place early or late along the sensory streams **(^40^Bizley, *et al.*, 2016)**. There are clear signs of multisensory responses in relatively early-or mid-stage of sensory areas, thus supporting the early theory **(^41^Gu, 2018)**. Our results are more consistent with the late-convergence theory in which multisensory momentary evidence are combined across modalities and time in decision areas such as LIP. However, this dichotomy between early and late theories does not necessarily make sense given the recurrent nature of the cortical circuitry. In a highly recurrent network, it is notoriously difficult to identify a node as a primary site of integration. Thus, integration might take place simultaneously across multiple sites but in such a way that the output of the computation is consistent across sites. For example, **Deneve, *et al.* ^42^** demonstrated how this could take place in a large recurrent network performing optimal multisensory integration, though their work did not consider the problem of temporal integration.

It might be possible to gain further insight into the distributed nature of multisensory decision making by combining the previous models with the one we have presented here. Such an extended model might explain why vestibular momentary evidence is tuned to velocity by the time they appear in MSTd **(^37^Laurens, *et al.*, 2017; ^41^Gu, 2018)**, and why this velocity tuned vestibular input does not appear to be integrated in LIP. It could also shed light on recent physiological experiments in which electrical microstimulation and chemical inactivation in MSTd could dramatically affect heading discrimination based on optic flow while this effect was largely negligible in the vestibular condition **(^43^Gu, *et al.*, 2012)**. By contrast, and in accord with our finding that LIP integrates vestibular acceleration, inactivating the vestibular cortex PIVC, where vestibular momentary evidence is dominated by acceleration **(^34^Chen, *et al.*, 2011a; ^37^Laurens, *et al.*, 2017)**, substantially diminished the macaque’s heading ability based on vestibular cue **(^44^Chen, *et al.*, 2016)**. Note, however, a detailed construction of such a model lies beyond the scope of the present study but will eventually be required for a multi-area theory of multisensory decision making.

### Computational models for multisensory decision making

Our results indicate that, at the population level, LIP implements an optimal solution for multisensory decision making under the assumption that the sensory inputs are encoded with ilPPC. This assumption is not perfectly satisfied in our experiment since the visual and vestibular inputs deviate from pure ilPPCs, but we saw that this deviation introduces only a minor information loss. While these results provide the first experimental support for the ilPPC theory of multisensory decision making, it will be important to test in future experiments other predictions of this framework. In particular, the ilPPC theory predicts that LIP activity encodes a full probability distribution over choices given the evidence so far **(^9^Beck, *et al.*, 2008)**. Testing this prediction thoroughly requires simultaneous recording of LIP ensemble, manipulating the cue reliability (motion profile or visual coherence) on a trial-by-trial basis, and preferably engaging the animals in a reaction-time task, all of which should be addressed in future studies.

There are of course other models of decision making which could potentially account for the responses we have observed in LIP **(^45^Chandrasekaran, 2017)**. In particular, it has been argued that LIP is part of a network of areas implementing point attractor networks **(^46^Wong and Wang, 2006; ^47^Wang, 2002)**. However, it is not immediately clear how this approach can be generalized to the type of decision we have considered here. Indeed, as we have seen, the optimal solution depends critically on the code that is used to encode the momentary evidence. To the extent that this code is close to an ilPPC, the optimal solution is to sum the inputs spikes, in which case one needs a line attractor network, which is effectively what our network approximates. Therefore, as long as these previous models of decision making are fine-tuned to approximate line attractor networks, and as long as they are fed ilPPCs as inputs, the two classes of models would be equivalent.

Training recurrent neural network (RNNs) on our task **(^48^Mante, *et al.*, 2013; ^49^Song, *et al.*, 2017)** provides a third alternative for modeling multisensory decision making. We also tried this approach and found that the resulting network was capable of reproducing the behavioral thresholds of the animal while exhibiting a wide variety of single neuron responses similar to what we saw in LIP (**Supplementary Figure 10**). Nonetheless, this approach has one major drawback: it makes it very difficult to understand how the network solves the task. We could try to reverse engineer the network, but given that an analytical solution can be derived from first principles for our task, and given that this solution is close to what we observed in LIP, it is unclear what insight could be gained from the recurrent network. In contrast, our ilPPC model provides a close approximation to the optimal solution, consistent with the experimental results, along with a clear understanding as to why this approach is optimal.

## Methods

### Subjects and Apparatus

All animal procedures were approved by the Animal Care Committee of Shanghai Institutes for Biological Sciences, Chinese Academy of Sciences and have been described previously in detail **(^12^Gu, *et al.*, 2008; ^19^Gu, *et al.*, 2006)**. Briefly, two male adult rhesus monkeys, Monkey P and Monkey M, weighing ∼8 kg, were chronically implanted with a lightweight plastic ring for head restraint and a scleral coil for monitoring eye movements (Riverbend Instruments). During experiments, the monkey sat comfortably in a primate chair mounted on top of a custom-built virtual reality system, which consisted of a motion platform (MOOG MB-E-6DOF/12/1000KG) and an LCD screen (∼30 cm of view distance and ∼90^°^ × 90^°^ of visual angle; HP LD4201), presenting vestibular and visual motion stimuli to the monkey, respectively. The stimuli were controlled by customized C++ software and synchronized with the electrophysiological recording system by TEMPO (Reflective Computing, U.S.A).

To tune the synchronization between vestibular and visual stimuli, we rendered a virtual world-fixed crosshair on the screen while projected a second crosshair at the same place on the screen using a real world-fixed laser pen. When the platform was moving, we carefully adjusted a delay parameter in the C++ software (with 1 ms resolution) until the two crosshairs aligned precisely together all the time, as verified by a high-speed camera (Meizu Pro 5) and/or a pair of back-to-back mounted photodiodes. This synchronization procedure was repeated occasionally over the whole period of data collection.

### Behavioral Tasks

#### Memory-guided Saccade Task

We used the standard memory-guided saccade task **(^50^Barash, *et al.*, 1991)** to characterize and select LIP cells for recording in the main decision-making experiments. The monkey fixated at a central fixation point for 100 ms and then a saccade target flashed briefly (500 ms) in the periphery. The monkey was required to maintain fixation during the delay period (1000 ms) until the fixation point extinguished and then saccade to the remembered target location within 1000 ms for a liquid reward. For all tasks in the present study, at any time when there existed a fixation point, trials were aborted immediately if the monkey’s gaze deviated from a 2^°^ × 2^°^ electronic window around the fixation point.

#### Multisensory Heading Discrimination Task

In the main experiments, we trained the monkeys to report their direction of self-motion in a two-alternative forced-choice heading discrimination task **(^12^Gu, *et al.*, 2008)** (**Figure 1**). The monkey initiated a trial by fixating on a central, head-fixed fixation point, and two choice targets then appeared. The locations of the two targets were determined case-by-case for each recording session (see below). After fixating for a short delay (100 ms), the monkey then began to experience a fixed-duration (1.5 s) forward motion in the horizontal plane with a small leftward or rightward component relative to straight ahead. The animals were required to maintain fixation during the presentation of the motion stimuli. At the end of the trial, the motion ended, and the monkey was required to maintain fixation for another 300–600 ms random delay (uniformly distributed) until the fixation point disappeared, at which point the monkey was allowed to make a saccade choice toward one of the two targets to report his perceived heading direction (left or right).

Across trials, nine heading angles (±8^°^, ±4^°^, ±2^°^, ±1^°^, and 0^°^) and three cue conditions (vestibular, visual, and combined) were jointly interleaved, resulting in 27 unique stimulus conditions, each of which was repeated 15 ± 3 (median ± m.a.d.) times per one session. In a vestibular or a visual trial, heading information was solely provided by inertial motion (real movement of the motion platform) or optic flow (simulated movement through a star field on the display), respectively, whereas in a combined trial, congruent vestibular and visual cues were provided synchronously. To maximize the behavioral benefit of cue integration, we balanced the monkey’s performance under the vestibular and the visual conditions by manipulating the motion coherence of the optic flow (the percentage of dots that moved coherently). The visual coherence was 12% and 8% for monkey P and M, respectively.

To ensure that the reliabilities of sensory cues varied throughout each trial, we used Gaussian-shape, rather than constant, velocity profiles for all motion stimuli. In the main experiments, the Gaussian profile had a displacement *d* = 0.2 *m* and a standard deviation *σ* = 210 *ms* (half duration at about 60% of the peak velocity), resulting in a peak velocity *v*_*max*_ = 0.37. *m/s* and a peak acceleration *a*_*max*_ = 1.1 *m/s*^2^ In the experiment where we sought to independently vary the peak times of velocity and acceleration (**Figure 3**), two additional sets of motion parameters were used. For the narrow-speed profile, *d* = 0.10 *m σ* = 150 *ms*, *v*_*max*_ = 0.37*m/s*, and *a*_*max*_ = 1.1 *m/s*^2^; for the wide-speed profile, *d* = 0.25 *m σ* =330 *ms*, *v*_*max*_ = 0.31 *m/s*, and *a*_*max*_ = 1.1 *m/s*^2^.

### Electrophysiology

We carried out extracellular single-unit recordings as described previously **(^12^Gu, *et al.*, 2008)** from four hemispheres in two monkeys. For each hemisphere, reliable area mapping was first achieved through cross-validation between structural MRI data and electrophysiological properties, including transition patterns of gray/white matter along each penetration, sizes of visual receptive/response field, strengths of spatial tuning to visual and vestibular heading stimuli, and activities in the memory-guided saccade task. Based on the mapping results, Area LIP was registered by its spatial relationships with other adjacent areas (VIP, Area 5, MSTd, etc.), its weak sensory encoding of heading information, and its overall strong saccade-related activity (**Supplementary Figure 1**). Our recording sites located in the ventral division of LIP, extending from 7–13 mm lateral to the midline and −5 mm (posterior) to +3 mm (anterior) relative to the interaural plane.

Once we encountered a well-isolated single unit in LIP, we first explored its response field (RF) by hand (using a flashing patch) and then examined its electrophysiological properties using the memory-guided saccade task. The saccade target in each trial was randomly positioned at one of the 8 locations 45^°^ apart on a circle centered on the fixation point (5^°^–25^°^ radius, optimized according to the cell’s RF location). We calculated online the memory-saccade spatial tuning for three response epochs: (1) visual response period, 75–400 ms from target onset; (2) delay period, 25–900 ms from target offset; and (3) presaccadic period, 200–50 ms before the saccade onset (**Supplementary Figure 2**). The cell’s spatio-temporal tunings were used to refine its RF location (via vector sum) and to determine its inclusion in the subsequent decision-making task. Since the decision-related activity of LIP neurons cannot be strongly predicted by the persistent activity during the delay period alone **(^26^Meister, *et al.*, 2013)** (**Supplementary Figure 4b**), we adopted a wider cell selection criterion than conventionally used, in which we included cells that have significant spatial selectivity for *any* of the three response epochs **(^26^Meister, *et al.*, 2013)** (one-way ANOVA, p < 0.05, 3–5 repetitions). If the cell met this criterion, then we recorded its decision-related activity while engaging the monkey in the main multisensory decision-making task, with the two choice targets being positioned in its RF and 180^°^ opposite to its RF, respectively.

Although we collected data from a relatively broad sample of LIP neurons, we nevertheless had two sampling biases during this process. First, we were biased toward cells with strong persistent activity so that our multisensory data could be better compared with previous unisensory data in the decision-making literature, where in most cases only these cells were recorded. Second, we were biased toward cells with RF close to the horizontal line through the fixation point. Unlike the classical random dot stimuli whose motion direction on the fronto-parallel plane can be aligned with the cell’s RF (and the choice targets) session by session, our self-motion stimuli were always on the horizontal plane and thus were not adjustable according to the cell’s RF on the fronto-parallel plane. As a result, the subjects had to make an additional mapping from their perceived heading directions (always left or right) to the choice targets (often inclined, and in extreme cases, up or down). Therefore, to make the task more intuitive to the monkeys and to minimize the potential influence of this mapping step on neural activity, we discarded a cell if the angle between the horizontal line and the line connecting the fixation point to its RF exceeded 60^°^, although we observed little change in monkeys’ behavior even when this angle approached 80^°^.

### Data Analysis

#### Psychophysics

To quantify the behavioral performance for both the monkeys and the model in the multisensory decision-making task, we constructed psychometric curves by plotting the proportion of “rightward” choices as a function of heading (**Figure 1c**) and fitted them with cumulative Gaussian functions **(^12^Gu, *et al.*, 2008)**. The psychophysical threshold for each cue condition was defined as the standard deviation of their respective Gaussian fit. The Bayesian optimal prediction of psychophysical threshold under the combined condition. *σ*_*prediction*_ was solved from the inversevariance rule **(^24^Knill and Richards,1996)**

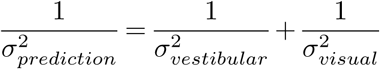

where *σ*_*vestibular*_ and *σ*_*visual*_ represent psychophysical thresholds under the vestibular and visual conditions, respectively.

#### Choice-related neural activities

We constructed peri-stimulus time histograms (PSTHs) for two epochs of interest in a trial, the decision formation epoch and the saccade epoch, by aligning raw spike trains to the stimulus onset and the saccade onset, respectively. Firing rates were computed in non-overlapping 10-ms bins and smoothed over time by convolving with a Gaussian kernel (*σ* = 50 *ms*). Unless otherwise noted, only correct trials were used in the following analyses, except for the ambiguous 0^°^ heading where we included all complete trials.

To illustrate the choice-related activity of a cell, we grouped the trials according to the monkey’s choice, i.e., trials ending up with a saccade toward the cell’s RF (IN trials) versus trials ending up with a saccade away from the cell’s RF (OUT trials), and computed the averaged PSTHs of these two groups of trials for each cue condition (**Figure 2a**). When averaged across cells, each cell’s PSTHs were normalized such that the cell’s overall firing rate had a dynamic range of [0, 1] (**Figure** 3). To quantify the strength of choice signals and better visualize ramping activities, we calculated choice divergence **(^23^Raposo, *et al.*, 2014)** for each 10-ms time bin and for each cue condition using receiver operating curve (ROC) analysis (**Figure 2b**). Choice divergence ranged from −1 to 1 and was defined as 2 × (AUC −0.5), where AUC represents the area under the ROC curve derived from PSTHs of IN and OUT trials. To capture the onset of choice signals, we computed a divergence time defined as the time of the first occurrence of a 250-ms window (25 successive 10-ms bins) in which choice divergence was consistently and significantly larger than 0 (**Figure 3c, f**). The statistical significance of choice divergence (p < 0.05, relative to the chance level of 0) was assessed by two-sided permutation test (1000 permutations). We also calculated a grand choice divergence which ignored temporal information and used all the spikes in the decision formation epoch (0–1500 ms from the stimulus onset). The same permutation test was performed on the grand choice divergence to determine whether a cell had overall significant choice signals for a certain cue condition (for example, in **Figure 2c**).

#### Linear Fitting of Mean Firing Rates

We fitted a linear weighted summation model to predict neural responses under the combined condition with those under the single cue conditions, using **(^12^Gu, *et al.,* 2008)**

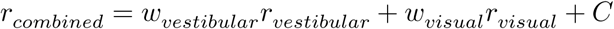

where *C* is a constant, and *r*_*combined*_, *r*_*vestibular*_, and *r*_*visual*_ are mean firing rates across a trial (0– 1500 ms from stimulus onset) for the three cue conditions, respectively. The weights for single cue conditions, *w*_*vestibular*_ and *w*_*visual*_, were determined by the least-squares method and plotted against each other to evaluate the heterogeneity of choice signals in the population for both LIP data and the model (**Supplementary Figure 7d**).

#### Fisher Information Analysis

To compute Fisher information **(^32^Seung and Sompolinsky, 1993)**, the full covariance matrix of the population responses is needed, but this requires simultaneously recording from hundreds of neurons, which is not accessible to us yet. Instead, we calculated the shuffled Fisher information, which corresponds to the information in a population of neurons in which correlations have been removed (typically via shuffling across trials, hence the name). Shuffled Fisher information is given by **(^33^Series, et al., 2004; ^51^Gu, et al., 2010)**:

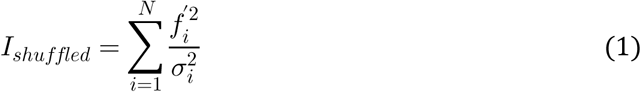

where *N* is the number of nein the population; for the *i*th neuron, 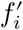 denotes the derivative of its local tuning curve, and 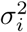 denotes the averaged variance of its responses around 0*°* heading. The tuning curve *f*_*i*_ was constructed from both correct and wrong trials grouped by heading angles, using spike counts in 250-ms sliding windows (advancing in 10-ms steps), and its derivative 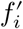 was obtained from the slope of a linear fit of *f*_*i*_ against headings. The variance 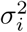 was computed for each heading angle and then averaged. To estimate the standard errors of *I*_*shuffled*_, we used a bootstrap procedure in which random samples of neurons were drawn from the population by resampling with replacement (1000 iterations). To compare the experimental data with the model, we repeated all the above steps on artificial LIP neurons in the model M2 and M3 (see below), with the inter-neuronal noise correlation being ignored as well (**Figure 6**).

Two caveats are noteworthy when interpreting the Fisher information results. First, since the slope of tuning curve *f ′* is squared in the right-hand side of equation (1), the Fisher information will always be non-negative regardless of the sign of *f ′* As a result, even when the motion speed was zero at the beginning of a trial, the population Fisher information already had a positive value because of the noisy tuning curves during that period. Second, since we ignored inter-neuronal noise correlations, *I*_*shuffled*_ is most likely very different from the true Fisher information and thus its value is arbitrary **(^33^Series, et al., 2004)**. Nonetheless, if we assume the noise correlation structure of LIP population is similar across cue conditions, we can still rely on the qualitative temporal evolution of *I*_*shuffled*_ to appreciate how multisensory signals are accumulated across time and cues in LIP.

### Network Simulation of ilPPC Framework

#### The responses of visual and vestibular neurons closely approximate ilPPC

As mentioned previously, an important assumption of ilPPC is that the amplitude of the sensory tuning curves be proportional to the nuisance parameters (in our case visual speed and vestibular acceleration) **(^9^Beck, et al., 2008)**. To check whether this is the case for the visual neurons, we analyzed the spatio-temporal tuning curves of neurons in area MSTd (data from **(^19^Gu, et al., 2006)**). We noticed that, for some neurons, the average tuning curves are not fully consistent with the ilPPC assumption (**Supplementary Figure 6a**). Briefly, the mean firing rate of an MSTd neuron at time *t* in response to a visual stimulus with heading θ can be well captured by

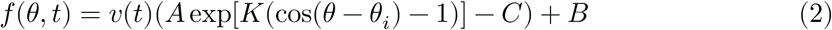

where *θ*_*i*_ denotes the preferred heading of the neuron *i* and *v*(*t*) is the velocity profile; *A*,*K*,*C,* and *B* correspond to the amplitude, the width, the null inhibition, and the baseline of its tuning curve, respectively. The ilPPC framework requires the *v*(*t*) term to be separable, namely, *f*(θ,*t*) = *h*(θ) *g*(*v*(*t*)), where *h*(θ) is a pure spatial component and *g*(*v*(*t*)) is a multiplicative gain function **(^9^Beck, et al., 2008; ^10^Ma, et al., 2006)**. In equation (2), this requirement is equivalent to *C* = 0 and *B* = 0, however, we found that some MSTd neurons often had non-zero baselines (*C* >0 and *B* > 0). This will be harmful to the optimality of the ilPPC framework because, for example, when (and thus the sensory reliability is zero), MSTd neurons still tend to generate background spikes, which will bring nothing but noise into the simply summed population activity of downstream areas in an ilPPC network.

To estimate the information loss due to this deviation, we simulated a population of MSTd neurons with heterogeneous spatio-temporal tuning curves similar to what has been found experimentally **(^19^Gu, et al., 2006)**. We calculated the information that can be decoded from the population by a series of optimal decoders *I*_*optimal*_ and that can be recovered by the ilPPC solution *I*_*ilppc*_. We assumed that the population responses in MSTd contains differential correlations **(^28^Moreno-Bote, et al., 2014)** such that the discrimination threshold of an ideal observer of MSTd activity was of the same order as the animal’s performance. Under such conditions, we found that the information loss (*I*_*optimal*_ – *I*_*ilppc*_)/*I*_*optimal*_ was around 5%. Detailed calculations of information loss are provided in the **Supplementary Materials**. Therefore, the population response of MSTd neurons provide a close approximation to an ilPPC, in the sense that simply summing the activity of MSTd neurons over time preserve 95% of the information conveyed by these neurons.

We also checked whether the ilPPC assumption holds in the case of vestibular neurons. Equation (2) above still provides a good approximation to vestibular tuning curves, except that is close to zero for most neurons **(^37^Laurens, et al., 2017)**, in which case the information less is even less pronounced.

#### Network Model Implementing the ilPPC solution (Model M1)

We extended a previous ilPPC network model for unisensory decision making **(^9^Beck, et al., 2008)** to our multisensory decision-making task. Two sensory layers, the vestibular layer and the visual layer, contained 100 linear-nonlinear-Poisson (LNP) neurons with bell-shape tuning curves to the heading direction (equation (2)). For the th neuron in the vestibular or visual layer, the probability of firing a spike at time step [*t*_*n*_ - δ*t*, *t*_*n*_] was given by

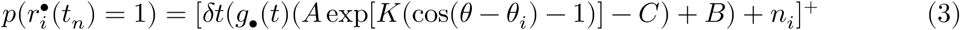

where • ϵ {VEST, VIS}, *A*,*K*,*C,* θ, and θ_*i*_ have the same meaning as in equation (2), *n*_*i*_ is a correlated noise term, and []+is the threshold-linear operator: [*x*]+ = max (*x*,0) The spatial tuning was gain-modulated by a time-dependent function *g* •(*t*), which modeled the reliability of the sensory evidence and took the form

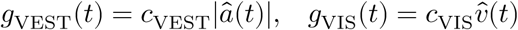

in which 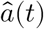 and 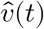 are the same acceleration and velocity profiles as the experiments but with the maximum values normalized to 1, respectively, whereas *c*VEST and *c*VIS are scaling parameters used to control the signal-to-noise ratio of sensory inputs and to balance the behavior performance between the two cue conditions like in the experiments. The noise *n*_*i*_ in equation (3) was generated by convolving independent Gaussian noise with a circular Gaussian kernel,

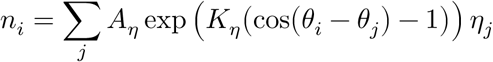

where η_*j*_. ∼ *i i d N* (0,1), and *A*η and *A*η were set to 10^−5^ and 2, respectively. Other parameters we used were: *A*= 60 Hz, *K* = 1.5, *C* = 10 Hz,*B* = 20 Hz, *c* VEST = *c* VIS = 2.4, δ*t* = 1 ms. Note that in equation (3), the gain *g* •(*t*) cannot be factored out because,B which is the same case as in MSTd (equation (2)). Accordingly, the neural code of M1’s sensory layers is not exact ilPPC **(^9^Beck, et al., 2008)**. However, it is still a close approximation to ilPPC, since we have shown in the previous section that MSTd is 95% ilPPC-compatible.

The two sensory layers then projected to 100 LNP neurons in the integrator layer. We distinguished the integrator layer from the LIP layer because there are reasons to believe that LIP reflects the integration of the evidence but may not implement the integration *per se* **(^27^Katz, et al., 2016)**. The integrator layer summed the sensory responses across both cues and time,

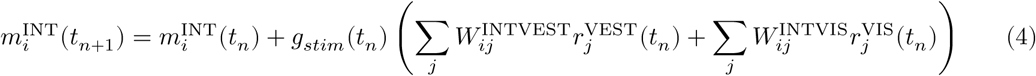

where 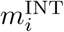 denotes the membrane potential proxy of neuron *i*, 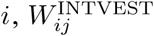 and 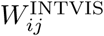 are matrices for the feedforward weights from the vestibular and visual layer to the integrator layer, respectively, and *g*_*stim*_ (*t*_*n*_) is an attentional gain factor (see below). Note that we ignored the issue of how neural circuits perform perfect integration and just assumed that they do. We could have simulated one of the known circuit solutions to this problem **(^52^Goldman, 2009)**, but this would not have affected our results, while making the simulation considerably more complicated.

The feedforward connections 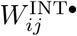 map the negative and positive heading directions onto the two saccade targets, i.e., neurons preferring −90^°^ and +90^°^ in the integrator layer, respectively, by

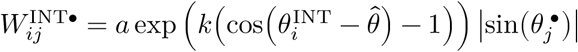

in which a step function 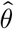 controls the mapping,

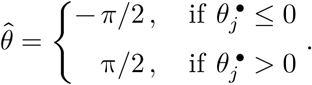

We used *a* = 20 and *k* = 4 in our simulations. After the linear step, the membrane potential proxy was used to determine the probability of the *ii*th integrator neuron firing a spike between times *t*_*n*_ and *t*_*n*_ + δ *t,*

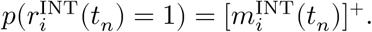

Finally, the LIP layer received excitatory inputs from the integrator layer, together with visual inputs triggered by the two saccade targets (sent from the target layer). In addition, there were also lateral connections in LIP to prevent saturation. In the linear step, the membrane potential proxy of the *i*th LIP neuron followed

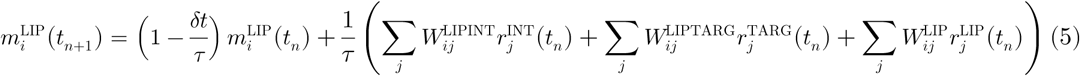

where the time constant, τ, was set to 100 ms; 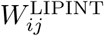 and 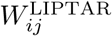 are weight matrices for the feedforward connections from the integrator layer and the target layer to the LIP layer, respectively, and 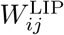 is the matrix for the recurrent connections within LIP. We used translation-invariant weights for all these connections,

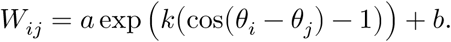

For 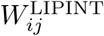, we used *a* = 15, *k* = 10, *b* = −3.6; for 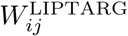, we used *a* = 8, *k* = 5, *b* = 0; and for 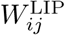, we used. *a* = 5, *k* = 10, *b* = −3; The term 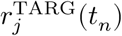 in equation (5) denotes the visual response of the *j*th neuron in the target layer induced by the two saccade targets,

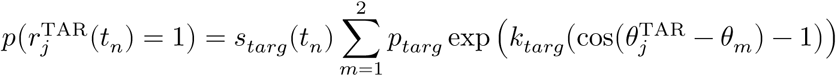

where θ1 = -π/2 and θ2 = -π/2, *p*_*targ*_= 0.050, and *k*_*targ*_ = 4. The term *s*_*targ*_ (*tn*) modeled the saliency of the targets: *s*_*targ*_ (*tn*) = 1 before stimulus onset and *s*_*targ*_ (*tn*) = 0.6 afterwards.

After the linear step done in equation (5), the probability of observing a spike from the th LIP neuron for the next time step was given by, again,

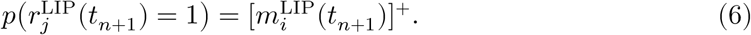

#### Decision Bound and Action Selection

To let the model make decisions, we endowed it with a stopping bound such that the evidence integration terminated when the peak activity in the LIP layer reached a threshold value. This mechanism generates premature decisions in our fixed duration task, which have been observed in the previous experiments **(^29^Kiani, et al., 2008)** as well as ours (see the main text). Specifically, once the firing rate of any neuron in the LIP layer (determined from equation (6)) exceeded Θ ^•^ = 37 HZ for a vestibular or a visual trial and for a combined trial, we blocked the sensory inputs to the integrator layer by setting the gain factor in equation (4) to zero:

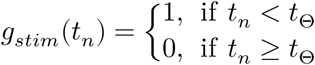

where *t*_Θ_ denotes the time of bound crossing. The instantaneous population activity at this time point *r*^LIP^(*t*_Θ_)was then used to determine the model’s choice, while the network dynamics continued to evolve until the end of the 1.5-s trial.

To read out the model’s choice, we trained a linear support vector machine (SVM) to classify the heading direction from *r*LIP(*t*_Θ_). We ran the network for 100 trials, used *r*LIP(*t*_Θ_) in 30 trials to train the SVM, and then applied the SVM on the remaining 70 trials to make decisions and generate psychometric functions of the model (with bootstrap 1000 times, **Figure 5a** and **Supplementary Figure 7a**). The SVM acts like (or even outperforms) a local optimal linear estimator (LOLE) trained by gradient descent **(^33^Series, et al., 2004)**. Importantly, such decoders could be implemented with population codes in a biologically realistic point attractor network tuned for optimal action selection in a discrimination task **(^53^Deneve, et al., 1999)**, which could correspond to downstream areas such as the motor layer of the superior colliculus **(^9^Beck, et al., 2008)**.

#### Heterogeneous ilPPC Network (M2)

In model M2, we generalized the homogeneous ilPPC network described above (model M1) to a heterogeneous one. Instead of taking perfect sums like in model M1, neurons in the integration layer of the model computed random linear combinations of vestibular and visual inputs. It is indeed been widely shown that integration weights *in vivo* are heterogeneous and are well-captured by “long-tailed” lognormal distributions (see for example **(^54^Song, et al., 2005)**). To simulate this in M2, we drew each synaptic weight wM2 in M2 from a lognormal distribution

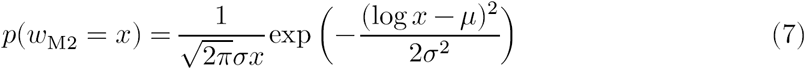

where *μ* and *σ* were chosen such that the expectation *u*(*w*_M2_) and the standard deviation *m*(*w*_M2_) of *w*_M2_ were both equal to its counterpart synaptic weight *w*M1 in model M1:

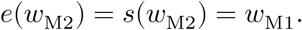

The parameters *μ* and *σ* in equation (7) were related to *σ* and *s* through

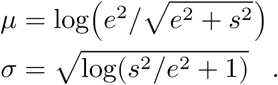

If *w*_M1_ < 0, a negative sign was added to the resulting *w*_M2_, since lognormal distributions are always non-negative.

#### Network with Short Integration Time Constant (M3)

We also simulated a sub-optimal model M3 in which the network does not integrate evidence over time. This was done by replacing equation (4) with

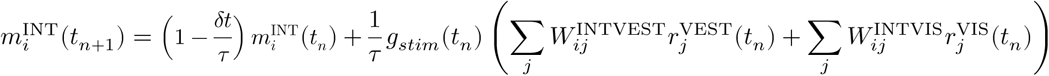

where τ = 100 *ms* and other terms are the same as in equation (4).

#### Linear Reproduction of M1 Response

To test whether the responses of the optimal and homogeneous model M1 can be linearly reproduced from responses of M2, M3, and the experimental data, we first calculated the “optimal traces” from M1 (**Figure 5b**), using

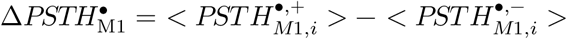

Wher **•** denotes three cue conditions (vestibular, visual, and combined),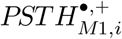 and 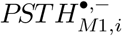 denote averaged PSTH for the *i*th LIP unit in the network M1 when the network makes correct choices towards the neuron’s preferred direction and null direction, respectively, and <·> denotes averaging across cells. To mimic the experimental procedure, only cells whose preferred directions were close to ±90*°* (with deviations less than 20^°^) were used. Similarly, we extracted single cell activities from M2, M3, the LIP data, and the MSTd data **(^19^Gu, et al., 2006)**

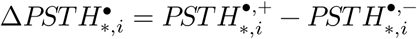

where * ∈ {M2, M3, LIP data, MSTd data}. Then we optimized sets of linear weights *w*_*_ to minimize the cost function

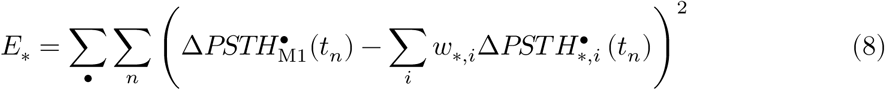

where, for example, *w*_LIP_,*i* represents the weight of the neuron in the LIP data when a downstream area reads out LIP dynamics linearly to reproduce the optimal traces. To reduce overfitting, we partitioned the data into two subsets along time by randomly assigning the time bins into two sets, one for fitting (*T*_fit_) and the other for validating (*T*_valid_). During fitting, when the validating error 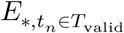 started increasing, we stopped the iteration, a procedure known as early stopping. The fitting results are shown in **Figure 5c–f**. Note that the Δ*PSTH* s in the cost function (equation (8)) grouped all the heading angles together. The results were qualitatively similar when the cost function included error terms calculated from each heading angle separately, i.e.,

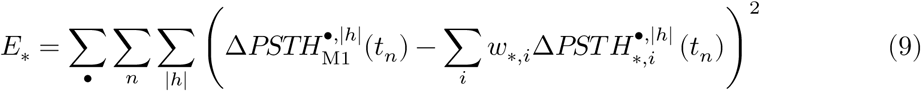

where |*h*| denotes the absolute value of heading angle (0^°^, 1^°^, 2^°^, 4^°^, 8^°^). The reconstructions of M1 traces with LIP activities using equation (9) are shown in **Supplementary Figure 9**.

To assess the robustness of the linear reconstruction, we randomly subsampled the same number of neurons (n = 50, without replacement) from the four data sets, performed the linear fitting, and repeated this procedure for 1000 times. The mean squared error and the distribution of readout weights of the fittings are shown in **Figure 5g, h**. To examine whether only a small fraction of cells contributed heavily to the fittings or whether the majority of cells did, we compared the distributions of weights from the four data sets with the distribution of weights from a random linear decoder. To do so, for each subsampling, we also generated a set of random readout weights from a rectified Gaussian distribution (**Figure 5h**, black curve) and computed the kurtosis of the distribution of weights from the random decoder as well as those from the four data sets (**Figure 5i**). The p-values were derived from the empirical subsampling distributions (two-tailed).

### Data and Code Availability

MATLAB code for the network model and the information loss calculation is available at the following public repository: https://github.com/hanhou/Multisensory-PPC. Experimental data and code for data analysis are available upon request to the authors.

## Supporting information

## Acknowledgments

We thank Jianyu Lu, Zhao Zeng, and Xuefei Yu for collecting part of the MSTd data, Wenyao Chen for monkey care and training, and Ying Liu for C++ software programming. This work was supported by grants from the National Natural Science Foundation of China Project (31761133014), the Strategic Priority Research Program of CAS (XDBS01070201), the Shanghai Municipal Science and Technology Major Project (2018SHZDZX05) to Y.G and by grants from the Simons Collaboration for the Global Brain and the Swiss National Science Foundation (#31003A_165831) to A.P.

## Author Contributions

H.H. and Y.G. conceived the project and designed the experiments. H.H., Q.Z., and Y.Z. performed the experiments. H.H. analyzed the data. H.H. and A.P. developed the models and implemented the simulations. H.H., A.P., and Y.G. wrote the manuscript.

## Supplementary Materials

**Supplementary Figure 1. Recording sites and reliable area mapping**

**Supplementary Figure 2. More example LIP cells**

**Supplementary Figure 3. Task-difficulty dependence of choice signals**

**Supplementary Figure 4. Macaque LIP is category-free**

**Supplementary Figure 5. De-mixing of choice and modality signals**

**Supplementary Figure 6. Information loss of ilPPC solution with heterogeneous MSTd population**

**Supplementary Figure 7. Model M2 achieves optimal behavior with heterogeneous units**

**Supplementary Figure 8. Example units in linear reconstruction of M1**

**Supplementary Figure 9. Linear reconstruction of M1 with cost function calculated for separated heading angles**

**Supplementary Figure 10. A trained recurrent neural network (RNN) performing multisensory decision-making task**

## References

1. Ratcliff, R. A theory of memory retrieval. Psychological review 85, 59 (1978).

2. Ratcliff, R. & McKoon, G. The diffusion decision model: theory and data for two-choice decision tasks. Neural computation 20, 873–922 (2008).

3. Ratcliff, R. & Rouder, J.N. Modeling response times for two-choice decisions. Psychological science 9, 347–356 (1998).

4. Ratcliff, R. & Smith, P.L. A comparison of sequential sampling models for two-choice reaction time. Psychol Rev 111, 333–367 (2004).

5. Laming, D.R.J. Information theory of choice-reaction times. (1968).

6. Bogacz, R., Brown, E., Moehlis, J., Holmes, P. & Cohen, J.D. The physics of optimal decision making: a formal analysis of models of performance in two-alternative forced-choice tasks. Psychol Rev 113, 700–765 (2006).

7. Gold, J.I. & Shadlen, M.N. The neural basis of decision making. Annual review of neuroscience 30, 535–574 (2007).

8. Drugowitsch, J., DeAngelis, G.C., Klier, E.M., Angelaki, D.E. & Pouget, A. Optimal multisensory decision-making in a reaction-time task. Elife 3 (2014).

9. Beck, J.M., et al. Probabilistic population codes for Bayesian decision making. Neuron 60, 1142–1152 (2008).

10. Ma, W.J., Beck, J.M., Latham, P.E. & Pouget, A. Bayesian inference with probabilistic population codes. Nat Neurosci 9, 1432–1438 (2006).

11. Fetsch, C.R., Pouget, A., DeAngelis, G.C. & Angelaki, D.E. Neural correlates of reliability-based cue weighting during multisensory integration. Nat Neurosci 15, 146–154 (2012).

12. Gu, Y., Angelaki, D.E. & Deangelis, G.C. Neural correlates of multisensory cue integration in macaque MSTd. Nat Neurosci 11, 1201–1210 (2008).

13. Fetsch, C.R., DeAngelis, G.C. & Angelaki, D.E. Bridging the gap between theories of sensory cue integration and the physiology of multisensory neurons. Nature reviews. Neuroscience 14, 429–442 (2013).

14. Shadlen, M.N. & Newsome, W.T. Neural basis of a perceptual decision in the parietal cortex (area LIP) of the rhesus monkey. Journal of neurophysiology 86, 1916–1936 (2001).

15. Shadlen, M.N. & Newsome, W.T. Motion perception: seeing and deciding. Proceedings of the National Academy of Sciences of the United States of America 93, 628–633 (1996).

16. Huk, A.C., Katz, L.N. & Yates, J.L. The Role of the Lateral Intraparietal Area in (the Study of) Decision Making. Annual review of neuroscience 40, 349–372 (2017).

17. Roitman, J.D. & Shadlen, M.N. Response of neurons in the lateral intraparietal area during a combined visual discrimination reaction time task. The Journal of neuroscience: the official journal of the Society for Neuroscience 22, 9475–9489 (2002).

18. Boussaoud, D., Ungerleider, L.G. & Desimone, R. Pathways for Motion Analysis - Cortical Connections of the Medial Superior Temporal and Fundus of the Superior Temporal Visual Areas in the Macaque. J Comp Neurol 296, 462–495 (1990).

19. Gu, Y., Watkins, P.V., Angelaki, D.E. & DeAngelis, G.C. Visual and nonvisual contributions to three-dimensional heading selectivity in the medial superior temporal area. The Journal of neuroscience: the official journal of the Society for Neuroscience 26, 73–85 (2006).

20. Chen, A., DeAngelis, G.C. & Angelaki, D.E. Representation of vestibular and visual cues to self-motion in ventral intraparietal cortex. The Journal of neuroscience: the official journal of the Society for Neuroscience 31, 12036–12052 (2011c).

21. Chen, A., DeAngelis, G.C. & Angelaki, D.E. Functional Specializations of the Ventral Intraparietal Area for Multisensory Heading Discrimination. The Journal of Neuroscience 33, 3567–3581 (2013).

22. Nikbakht, N., Tafreshiha, A., Zoccolan, D. & Diamond, M.E. Supralinear and Supramodal Integration of Visual and Tactile Signals in Rats: Psychophysics and Neuronal Mechanisms. Neuron 97, 626-639.e628 (2018).

23. Raposo, D., Kaufman, M.T. & Churchland, A.K. A category-free neural population supports evolving demands during decision-making. Nat Neurosci 17, 1784–1792 (2014).

24. Knill, D.C. & Richards, W. Perception as Bayesian inference (Cambridge University Press, 1996).

25. Park, I.M., Meister, M.L., Huk, A.C. & Pillow, J.W. Encoding and decoding in parietal cortex during sensorimotor decision-making. Nat Neurosci 17, 1395–1403 (2014).

26. Meister, M.L., Hennig, J.A. & Huk, A.C. Signal multiplexing and single-neuron computations in lateral intraparietal area during decision-making. The Journal of Neuroscience 33, 2254–2267 (2013).

27. Katz, L.N., Yates, J.L., Pillow, J.W. & Huk, A.C. Dissociated functional significance of decision-related activity in the primate dorsal stream. Nature advance online publication (2016).

28. Moreno-Bote, R., et al. Information-limiting correlations. Nat Neurosci 17, 1410–1417 (2014).

29. Kiani, R., Hanks, T.D. & Shadlen, M.N. Bounded integration in parietal cortex underlies decisions even when viewing duration is dictated by the environment. The Journal of neuroscience: the official journal of the Society for Neuroscience 28, 3017–3029 (2008).

30. Scott, B.B., et al. Fronto-parietal Cortical Circuits Encode Accumulated Evidence with a Diversity of Timescales. Neuron 95, 385-398.e385 (2017).

31. Beck, J., Bejjanki, V.R. & Pouget, A. Insights from a simple expression for linear fisher information in a recurrently connected population of spiking neurons. Neural computation 23, 1484–1502 (2011).

32. Seung, H.S. & Sompolinsky, H. Simple models for reading neuronal population codes. Proceedings of the National Academy of Sciences of the United States of America 90, 10749–10753 (1993).

33. Series, P., Latham, P.E. & Pouget, A. Tuning curve sharpening for orientation selectivity: coding efficiency and the impact of correlations. Nat Neurosci 7, 1129–1135 (2004).

34. Chen, A., DeAngelis, G.C. & Angelaki, D.E. A comparison of vestibular spatiotemporal tuning in macaque parietoinsular vestibular cortex, ventral intraparietal area, and medial superior temporal area. The Journal of neuroscience: the official journal of the Society for Neuroscience 31, 3082– 3094 (2011a).

35. Fetsch, C.R., et al. Spatiotemporal Properties of Vestibular Responses in Area MSTd. Journal of neurophysiology 104, 1506–1522 (2010).

36. Smith, A.T., Greenlee, M.W., DeAngelis, G.C. & Angelaki, D.E. Distributed Visual–Vestibular Processing in the Cerebral Cortex of Man and Macaque. Multisensory Research 30, 91–120 (2017).

37. Laurens, J., et al. Transformation of spatiotemporal dynamics in the macaque vestibular system from otolith afferents to cortex. Elife 6, e20787 (2017).

38. Lisberger, S.G. & Movshon, J.A. Visual motion analysis for pursuit eye movements in area MT of macaque monkeys. The Journal of neuroscience: the official journal of the Society for Neuroscience 19, 2224–2246 (1999).

39. Churchland, A.K., et al. Variance as a signature of neural computations during decision making. Neuron 69, 818–831 (2011).

40. Bizley, J.K., Jones, G.P. & Town, S.M. Where are multisensory signals combined for perceptual decision-making? Current opinion in neurobiology 40, 31–37 (2016).

41. Gu, Y. Vestibular signals in primate cortex for self-motion perception. Current opinion in neurobiology 52, 10–17 (2018).

42. Deneve, S., Latham, P.E. & Pouget, A. Efficient computation and cue integration with noisy population codes. Nat Neurosci 4, 826–831 (2001).

43. Gu, Y., Deangelis, G.C. & Angelaki, D.E. Causal links between dorsal medial superior temporal area neurons and multisensory heading perception. The Journal of neuroscience: the official journal of the Society for Neuroscience 32, 2299–2313 (2012).

44. Chen, A., Gu, Y., Liu, S., DeAngelis, G.C. & Angelaki, D.E. Evidence for a Causal Contribution of Macaque Vestibular, But Not Intraparietal, Cortex to Heading Perception. The Journal of neuroscience: the official journal of the Society for Neuroscience 36, 3789–3798 (2016).

45. Chandrasekaran, C. Computational principles and models of multisensory integration. Current opinion in neurobiology 43, 25–34 (2017).

46. Wong, K.F. & Wang, X.J. A recurrent network mechanism of time integration in perceptual decisions. The Journal of neuroscience: the official journal of the Society for Neuroscience 26, 1314–1328 (2006).

47. Wang, X.J. Probabilistic decision making by slow reverberation in cortical circuits. Neuron 36, 955–968 (2002).

48. Mante, V., Sussillo, D., Shenoy, K. & Newsome, W. Context-dependent computation by recurrent dynamics in prefrontal cortex. Nature 503, 78–84 (2013).

49. Song, H.F., Yang, G.R. & Wang, X.J. Reward-based training of recurrent neural networks for cognitive and value-based tasks. Elife 6 (2017).

50. Barash, S., Bracewell, R.M., Fogassi, L., Gnadt, J.W. & Andersen, R.A. Saccade-related activity in the lateral intraparietal area. I. Temporal properties; comparison with area 7a. Journal of neurophysiology 66, 1095–1108 (1991).

51. Gu, Y., Fetsch, C.R., Adeyemo, B., Deangelis, G.C. & Angelaki, D.E. Decoding of MSTd population activity accounts for variations in the precision of heading perception. Neuron 66, 596–609 (2010).

52. Goldman, M.S. Memory without Feedback in a Neural Network. Neuron 61, 621–634 (2009).

53. Deneve, S., Latham, P.E. & Pouget, A. Reading population codes: a neural implementation of ideal observers. Nat Neurosci 2, 740–745 (1999).

54. Song, S., Sjostrom, P.J., Reigl, M., Nelson, S. & Chklovskii, D.B. Highly nonrandom features of synaptic connectivity in local cortical circuits. PLoS biology 3, e68 (2005).

