## Supplementary material for "Neural Correlates of Optimal Multisensory Decision Making"

### Supplementary Modeling

##### Estimating the Information Loss of ilPPC Solution with MSTd-like Neural Population

As discussed in the main text, a prerequisite of the ilPPC solution being optimal is the spatio-temporal separability of sensory tuning curves. In practice, this requirement can be loosened by allowing a time-independent baseline to exist in the right-hand side of Equation 2, i.e., $B > 0$ but $C = 0$, since this baseline term can be readily removed without loss of information by, for example, a simple rectified linear unit (ReLU) layer. A recent study shows that the tuning curves of most vestibular neurons fall into this category (**[1] Laurens, *et al.*, 2017**). However, our analysis of the visual responses of MSTd neurons (n = 195, data from (**[2] Gu, *et al.*, 2006**)) suggests that this is not the case in MSTd (**Supplementary Figure 6a**, upper panel). The downward shift of firing rates around the non-preferred directions points to a time-dependent baseline component in the visual tunings, i.e., $B > 0$ and $C > 0$ in Equation 1, which cannot be simply eliminated with a layer of ReLU units. This implies that summing the activity of MSTd neurons over time will necessarily result in an information loss though the amplitude of this loss is unclear.

To estimate this information loss, we simulated a heterogeneous MSTd population with baseline-changing tuning curves and computed the proportion of encoded information that can be recovered by the ilPPC solution (i.e., a simple temporal integration). Specifically, we modeled the mean firing rate of neuron $i$ in response to a stimulus with heading $\theta$ at time $t$ by

$$f_i(\theta,t) = cv(t)(A_i\,exp[K_i(cos(\theta-\theta_{pref,i})-1)] - \beta_i B_i) + B_i,\;\; i = 1,2,\dots,N \qquad S\ 1$$

where $c \in [0,1]$ represents the motion coherence of the visual stimulus, $v(t)$ is the Gaussian velocity profile normalized to $[0,1]$, $A_i$, $K_i$, and $B_i$ controls the peak, the width, and the baseline of the tuning curve, respectively, and $\beta_i \in [0,1]$ controls how strong the baseline varies with time. The preferred headings, $\theta_{pref,i}$, were drawn uniformly from $(-\pi,\pi]$. To account for the heterogeneous tuning properties of real MSTd neurons, we sampled $A_i$, $B_i$, $\beta_i$, and the tuning width (full width at the half maximum, or $FWHM_i$) from Gamma distributions with the following mean and standard deviation values ($\beta_i$ larger than 1 were truncated to 1):

| | $A$ | $B$ | $FWHM$ | $\beta$ |
|---|---|---|---|---|
| Mean | 50 Hz | 20 Hz | 125° | 0.6 |
| Std. | 30 Hz | 20 Hz | 50° | 0.4 |

Then we found $K_i$ by $K_i = \frac{\log(0.5)}{\cos(FWHM_i/2-1\ )}$. These tuning parameters were chosen so that both the averaged and the single cell spatio-temporal tuning curves were similar to the MSTd data at the 100% visual coherence (**Supplementary Figure 6a**, lower panel). Note that we used 10% visual coherence ($c = 0.1$ in Equation S1) for the following calculations, a value close to what we used in the present study.

Next, we introduced correlated noise into this heterogeneous population via a stimulus- and time-dependent covariance matrix $\mathbf{\Sigma}(\theta, t)$,

$$\Sigma_{ij}(\theta, t) = c_{ij}\sqrt{f_i(\theta, t) f_j(\theta, t)} + \frac{\epsilon}{v(t)}\frac{\partial}{\partial\theta} f_i(\theta, t)\frac{\partial}{\partial\theta} f_j(\theta, t). \qquad S\ 2$$

The first term of the right-hand side is exponentially decaying pairwise correlations with a Fano factor = 1 and correlation coefficients

$$c_{ij} = (1-\rho)\delta_{ij} + \rho\, exp[\kappa_c(cos(\theta_{pref,i} - \theta_{pref,j}) - 1)] \qquad S\ 3$$

where we used $\rho = 0.1$ and $\kappa_c = 2$, and $\delta_{ij}$ is the Kronecker delta; the second term represents correlations proportional to the product of the derivatives of the tuning curves, referred to as differential correlation **([3] Moreno-Bote, *et al.*, 2014)**. The strength of the differential correlation is controlled by $\epsilon$, and the factor $1/v(t)$ is used to make the two terms of the right-hand side of Equation S2 have the same dependence on $v(t)$.

Having modeled both the tuning curves $\boldsymbol{f}(\theta, t)$ and the covariance matrix $\mathbf{\Sigma}(\theta, t)$, we set out to estimate the information. Without loss of generality, we considered information around $\theta = 0$ and we dropped the term $\theta$ hereafter for clarity. We focused on two kinds of information: 1, the amount of total information that can be decoded by a series of locally optimal linear estimators (LOLEs) optimized for each time $t$, denoted $I_{optimal}$, and 2, the amount of information that can be recovered by the ilPPC solution, that is, by a single LOLE acting on the linear summation of spikes over the entire trial, denoted $I_{ilPPC}$. Thus for $I_{optimal}$, by definition,

$$I_{optimal} = \int_0^T \dot{I}_{optimal}(t)dt \qquad S\ 4$$

where $T$ is the trial duration (2 s) and $\dot{I}_{optimal}(t)$ is the optimal information *rate* at time $t$. For LOLEs, $\dot{I}_{optimal}(t)$ is equal to the linear Fisher information **([4] Abbott and Dayan, 1999; [5] Beck, *et al.*, 2011; [3] Moreno-Bote, *et al.*, 2014)**,

$$\dot{I}_{optimal}(t) = \boldsymbol{f}'(t)^T \, \boldsymbol{\Sigma}^{-1}(t) \, \boldsymbol{f}'(t) \; . \qquad S\ 5$$

To obtain $\boldsymbol{f}'(t)$, we first computed the derivatives of tuning curves with respect to $\theta$ around $\theta = 0$ at time $t$, using Equation 1, and then multiplied the derivatives by a factor of 2. We did the second step because it has been shown that, in MSTd, the slopes of the local tuning curves are approximately two times larger than those derived from the global tuning curves **([6] Gu, *et al.*, 2010)**. Note, however, that this local sharpening of tuning curves will not affect the covariance $\boldsymbol{\Sigma}$, since the mean firing rate at $\theta = 0$ does not change. Combing $\boldsymbol{f}'(t)$ and $\boldsymbol{\Sigma}(t)$ in Equations S2-S3, we can thus calculate $I_{optimal}$ using Equations S4 and S5.

Similarly, for $I_{ilPPC}$, we have

$$I_{ilPPC} = \boldsymbol{f}'^{\,T}_{\boldsymbol{sum}} \, \boldsymbol{\Sigma}^{-1}_{\boldsymbol{sum}} \, \boldsymbol{f}'_{\boldsymbol{sum}} \; , \qquad S\ 6$$

except that here $\boldsymbol{f}_{\boldsymbol{sum}}$ and $\boldsymbol{\Sigma}_{\boldsymbol{sum}}$ are tuning curves and covariance matrix of another $N$ hypothetical LIP cells that implement the ilPPC solution via linear summation of spikes from their corresponding $N$ MSTd neurons. Therefore, by definition,

$$\boldsymbol{f}'_{\boldsymbol{sum}} = \frac{\partial}{\partial\theta} \int_0^T \boldsymbol{f}(t) dt = \int_0^T \boldsymbol{f}'(t) dt \, . \qquad S\ 7$$

To calculate $\boldsymbol{\Sigma}_{\mathbf{sum}}$, using the property of covariance

$$\Sigma_{sum,ij} = Cov\left(\int_0^T r_i(t)dt \, , \int_0^T r_j(t)dt\right) = \int_0^T \int_0^T dt_1 \; dt_2 \, Cov\left(r_i(t_1), r_j(t_2)\right) \qquad S\ 8$$

and noting that the firing rate of MSTd neurons $r(t)$ is assumed to be independent across time (see Discussion in the main text), namely,

$$Cov\left(r_i(t_1), r_j(t_2)\right) = \delta(t_1 - t_2) \; Cov\left(r_i(t_1), r_j(t_1)\right) = \delta(t_1 - t_2) \Sigma_{ij}(t_1), \qquad S\ 9$$

we have

$$\boldsymbol{\Sigma_{sum}} = \int_0^T \int_0^T \delta(t_1 - t_2)\boldsymbol{\Sigma}(t_1)dt_1 dt_2 = \int_0^T \boldsymbol{\Sigma}(t_1)dt_1 \int_0^T \delta(t_1 - t_2)dt_2 = \int_0^T \boldsymbol{\Sigma}(t)dt \ . \qquad S\ 10$$

Now we can calculate $I_{ilPPC}$ by pluging Equations S7 and S10 into Equation S6. In our simulations, we discretized the time into small steps of $\Delta t = 50$ ms, and the choice of $\Delta t$ did not influence our results much.

**Supplementary Figure 6b** shows $I_{optimal}$ and $I_{ilPPC}$ as a function of the number of neurons $N$ for two different $\epsilon$ s. When $\epsilon = 0$, the total optimal information $I_{optimal}$ did not saturate as $N$ went to infinity, and $I_{ilPPC}$, although smaller than $I_{optimal}$, also scaled almost linearly with $N$ (black solid and dashed curves). This is because for a heterogenous population, the first part of correlation in Equation S2 alone is not information-limiting **([7] Shamir and Sompolinsky, 2006; [8] Ecker*, et al.*, 2011; [3] Moreno-Bote*, et al.*, 2014)**. This regime however is unrealistic as it predicts that the information in MSTd would be considerably larger than the information available in the behavior of the animal. Indeed, single neurons often performs only slightly worse than the animal, suggesting that the information they convey is only slightly less than the information in the behavior. Since information is proportional to the number of neurons for $\epsilon = 0$, it follows that the information in a large neuronal population would vastly exceed the information in the behavior. Of course, downstream neurons may read out MSTd suboptimally, but [9] **Pitkow*, et al.* (2015)** have shown that choice correlations in MSTd are consistent with near optimal read out of MSTd activity.

Therefore, we assumed that there are significant information-limiting correlations, or equivalently, differential correlations **([3] Moreno-Bote*, et al.*, 2014)** in MSTd. When differential correlations were present ($\epsilon = 0.0015$), both $I_{optimal}$ and $I_{ilPPC}$ saturated rapidly with increasing $N$ (red solid and dashed lines in **Supplementary Figure 6b**). The upper limit at which they saturated (blue dashed curves in **Supplementary Figure 6b, d**) were predicted by

$$I_{optimal,\infty}(\epsilon) = \int_0^T \dot{I}_{optimal,\infty}(\epsilon)dt = \int_0^T \left(\frac{\epsilon}{v(t)}\right)^{-1} dt = \frac{1}{\epsilon}\int_0^T v(t)dt\,. \qquad S\ 11$$

Here we used the fact that for any total covariance matrix of the form

$$\boldsymbol{\Sigma} = \boldsymbol{\Sigma_0} + \epsilon \boldsymbol{f'}\boldsymbol{f'}^T \qquad S\ 12$$

where $\boldsymbol{\Sigma_0}$ does not limit information, the information when $N$ goes to infinity will be **([3] Moreno-Bote*, et al.*, 2014)**

$$\dot{I}_{\infty} = \lim_{\dot{I}_0 \to \infty} \left( \frac{\dot{I}_0}{1 + \epsilon \dot{I}_0} \right) = \frac{1}{\epsilon} \; , \qquad S\ 13$$

which depends only on $\epsilon$, where $\dot{I}_0$ is the non-saturating information corresponding to $\epsilon = 0$.

To assess how the baseline problem affects the optimality of ilPPC, we calculated the percentage of information preserved by the ilPPC solution,

$$Optimality\ ratio = 1 - InformationLoss\% = \frac{I_{ilPPC}}{I_{optimal}} \times 100\% \; . \qquad S\ 14$$

As shown in **Supplementary Figure 6c**, for $\epsilon = 0.0015$, the optimality ratio of ilPPC solution gradually increased with $N$ and reached 95% when $N = 10000$ (red curve). In contrast, for biologically implausible value of $\epsilon = 0$, the ilPPC solution is much more sensitive to the time-shifting baseline of MSTd tuning curves (> 40% information lost at $N = 10000$; black curve).

Obviously, the choice of $\epsilon$ is critical to our calculation, and the reason we have chosen $\epsilon = 0.0015$ is illustrated in **d**. We plotted the optimality and the psychophysical threshold $\sigma_{psy}$ together as a function of $\epsilon$. The threshold $\sigma_{psy}$ is defined as a small heading deviation around $\theta = 0$ that could be discriminated at 84% correct by an ideal observer **([10] Gu, *et al.*, 2011)**, which is directly comparable with the animal's psychophysical threshold reported in our experiments (around 4°). Thus, we have

$$\sigma_{psy} = \sqrt{2}\sigma_{LOLE} = \frac{\sqrt{2}}{\sqrt{I_{ilPPC}}} \; , \qquad S\ 15$$

where $\sigma_{LOLE}$ is the standard deviation of a LOLE, which is equal to the inverse of the linear Fisher information **([10] Gu, *et al.*, 2011; [3] Moreno-Bote, *et al.*, 2014)**. As predicted, when $\epsilon$ decreased, both $\sigma_{psy}$ and the optimality decreased, and we found that for a value around $\sigma_{psy} = 4°$, $\epsilon$ should be around 0.0015, and corresponding to an information loss around 5%.

Furthermore, we checked whether our results were sensitive to the distributions of the baselines $B_i$. As expected, the optimality of ilPPC decreased gradually from 100% to 84% as the averaged baseline increased from 0 Hz to 40 Hz (**Supplementary Figure 6e**). However, the gradient of the optimality around the biologically realistic regime (red star) was small enough for us to be confident about the stability of our estimates.

Taken together, we have shown that a moderate deviation from the ilPPC's prerequisite ($B_i = 0, \forall i$) will only result in ~5% information loss for the ilPPC solution.

### References


1. Laurens, J., *et al.* Transformation of spatiotemporal dynamics in the macaque vestibular system from otolith afferents to cortex. *Elife* **6**, e20787 (2017).

2. Gu, Y., Watkins, P.V., Angelaki, D.E. & DeAngelis, G.C. Visual and nonvisual contributions to three-dimensional heading selectivity in the medial superior temporal area. *The Journal of neuroscience : the official journal of the Society for Neuroscience* **26**, 73-85 (2006).

3. Moreno-Bote, R., *et al.* Information-limiting correlations. *Nat Neurosci* **17**, 1410-1417 (2014).

4. Abbott, L.F. & Dayan, P. The effect of correlated variability on the accuracy of a population code. *Neural computation* **11**, 91-101 (1999).

5. Beck, J., Bejjanki, V.R. & Pouget, A. Insights from a simple expression for linear fisher information in a recurrently connected population of spiking neurons. *Neural computation* **23**, 1484-1502 (2011).

6. Gu, Y., Fetsch, C.R., Adeyemo, B., Deangelis, G.C. & Angelaki, D.E. Decoding of MSTd population activity accounts for variations in the precision of heading perception. *Neuron* **66**, 596-609 (2010).

7. Shamir, M. & Sompolinsky, H. Implications of neuronal diversity on population coding. *Neural computation* **18**, 1951-1986 (2006).

8. Ecker, A.S., Berens, P., Tolias, A.S. & Bethge, M. The effect of noise correlations in populations of diversely tuned neurons. *The Journal of neuroscience : the official journal of the Society for Neuroscience* **31**, 14272-14283 (2011).

9. Pitkow, X., Liu, S., Angelaki, Dora E., DeAngelis, Gregory C. & Pouget, A. How Can Single Sensory Neurons Predict Behavior? *Neuron* **87**, 411-423 (2015).

10. Gu, Y., *et al.* Perceptual learning reduces interneuronal correlations in macaque visual cortex. *Neuron* **71**, 750-761 (2011).

11. Raposo, D., Kaufman, M.T. & Churchland, A.K. A category-free neural population supports evolving demands during decision-making. *Nat Neurosci* **17**, 1784-1792 (2014).

12. Meister, M.L., Hennig, J.A. & Huk, A.C. Signal multiplexing and single-neuron computations in lateral intraparietal area during decision-making. *The Journal of Neuroscience* **33**, 2254-2267 (2013).

13. Mante, V., Sussillo, D., Shenoy, K. & Newsome, W. Context-dependent computation by recurrent dynamics in prefrontal cortex. *Nature* **503**, 78-84 (2013).

14. Martens, J. Deep learning via Hessian-free optimization. in *Proceedings of the 27th International Conference on Machine Learning (ICML-11)* (2010).

15. Martens, J. & Sutskever, I. Learning recurrent neural networks with hessian-free optimization. in *Proceedings of the 28th International Conference on Machine Learning (ICML-11)* 1033-1040 (2011).

### Supplementary Figures

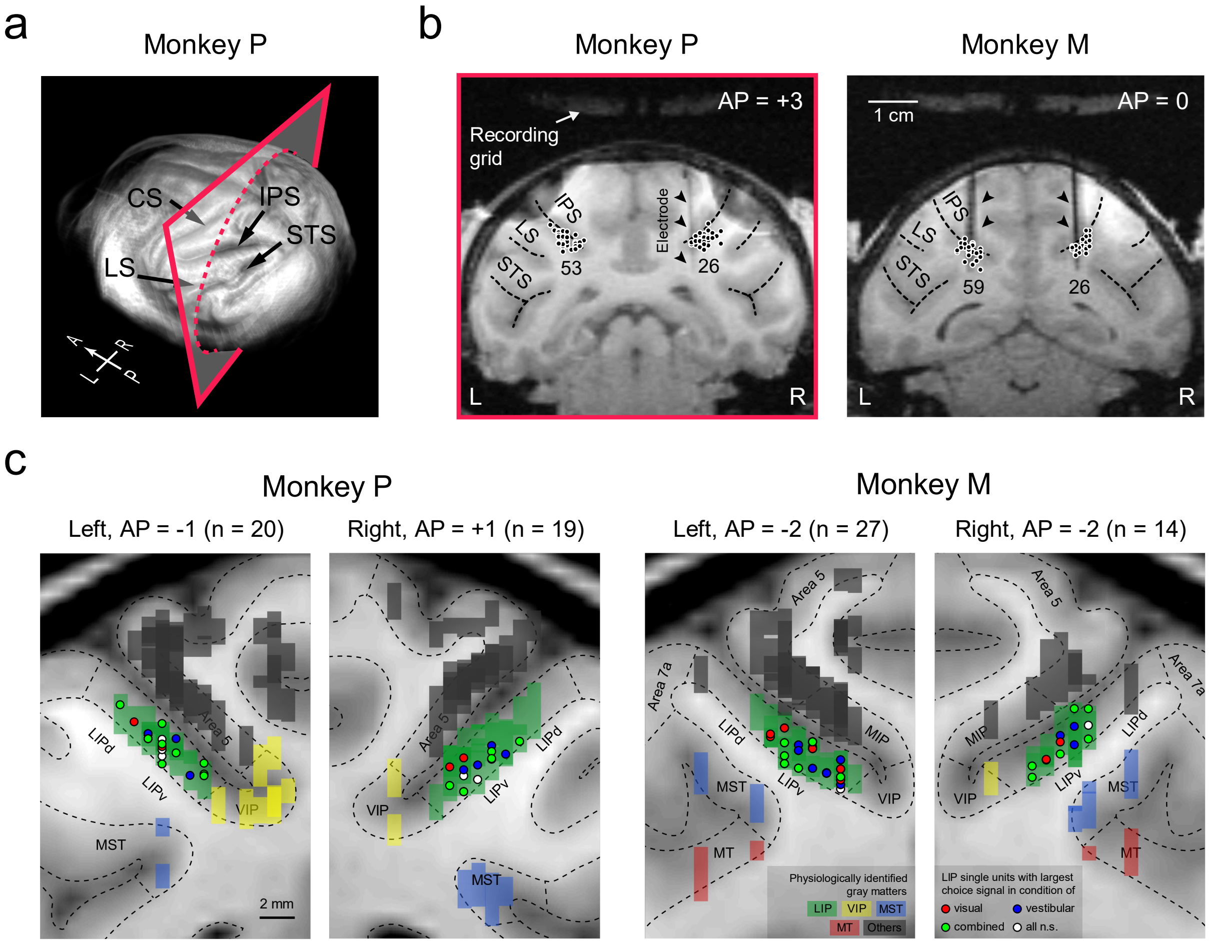


**Supplementary Figure 1. Recording sites and reliable area mapping**

**(a)** A 3-D reconstruction of MRI scanning result from Monkey P. The semi-transparent plane indicates the approximate location of the coronal section drawn in the left panel of **b**. IPS, intraparietal sulcus; STS, superior temporal sulcus; LS, lateral sulcus; CS, central sulcus; A, P, L, and R stand for anterior, posterior, left, and right, respectively.

**(b)** Representative coronal sections of the two monkeys. Recording sites are labeled by small dots, with the cell numbers for each hemisphere shown alongside. Vertical dark traces caused by artifacts of tungsten electrodes during MRI scanning (black arrowheads) were used as reference points for area mapping. Note that all the 164 cells are projected onto the two sections, making some units artificially fall outside the gray matters. AP, location (in mm) relative to the interaural plane along the anterior-posterior axis. Our recording sites extended from AP = -5 to +3.

**(c)** Four enlarged coronal sections demonstrating reliable area mapping results. MRI data (background) are superimposed by locations of gray matters (vertical colored rectangles) and single cells (colored circles). Gray matter's location and identity were registered by cross-validation between anatomical relationships and electrophysiological properties (see **Methods**). LIP (green), lateral intraparietal area; VIP (yellow), ventral intraparietal area; MST (blue), medial superior temporal area; MT (red), medial temporal area; Others (gray), other gray matters that were task-irrelevant. The color of each circle indicates the cue condition under which the corresponding cell has the strongest and significant choice signal (grand choice divergence, see **Methods**). Red, visual; blue, vestibular; green, combined; white, the cell does not have significant choice signal in any of the three conditions. On each section, cells located within 2 mm of that section are shown.

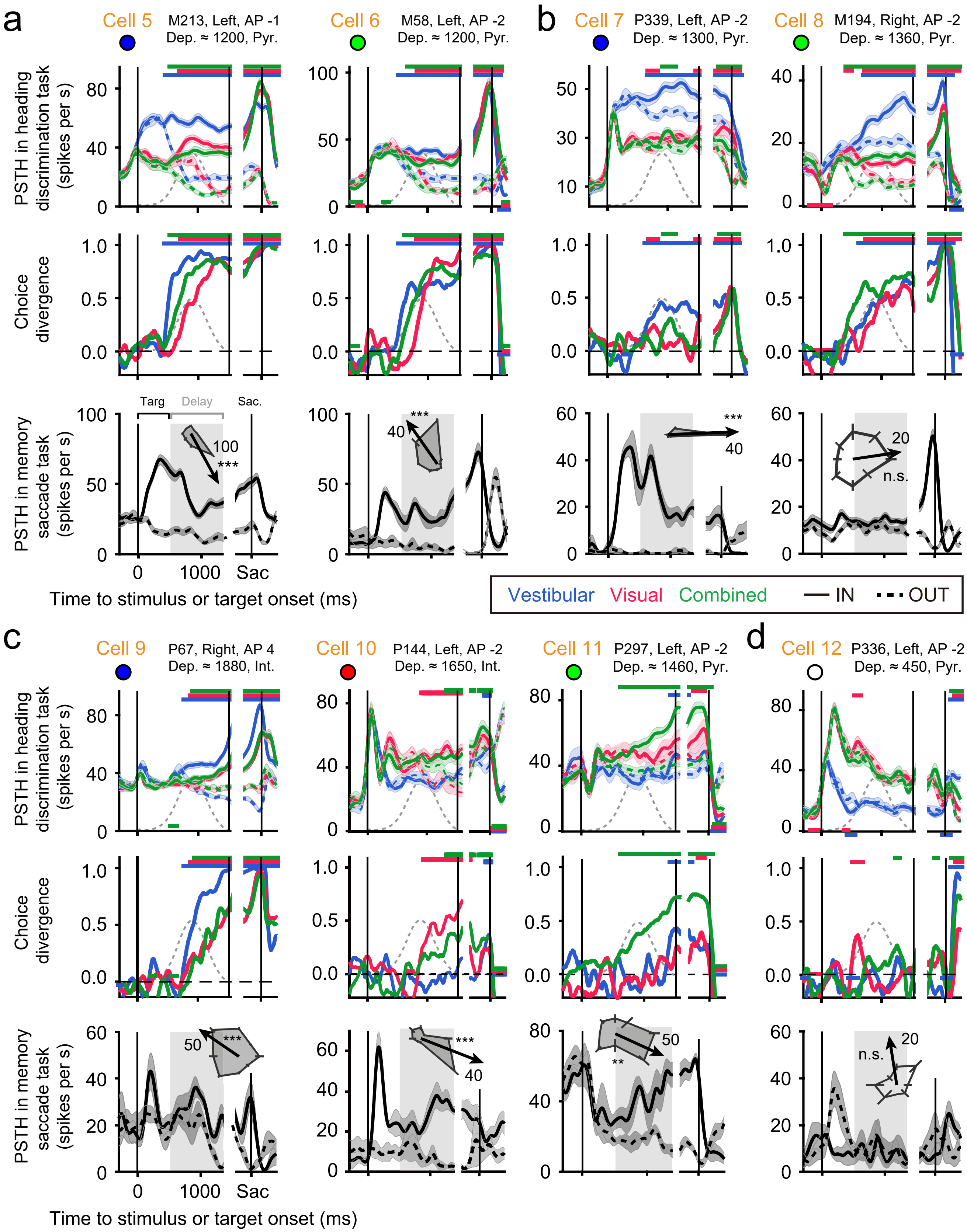
a
Cell 5 M213, Left, AP -1 Dep. ≈ 1200, Pyr.
Cell 6 M58, Left, AP -2 Dep. ≈ 1200, Pyr.
b
Cell 7 P339, Left, AP -2 Dep. ≈ 1300, Pyr.
Cell 8 M194, Right, AP -2 Dep. ≈ 1360, Pyr.
PSTH in heading discrimination task (spikes per s)
Choice divergence
PSTH in memory saccade task (spikes per s)
Targ
Delay
Sac.
Time to stimulus or target onset (ms)
Vestibular Visual Combined
IN
OUT
c
Cell 9 P67, Right, AP 4 Dep. ≈ 1880, Int.
Cell 10 P144, Left, AP -2 Dep. ≈ 1650, Int.
Cell 11 P297, Left, AP -2 Dep. ≈ 1460, Pyr.
d
Cell 12 P336, Left, AP -2 Dep. ≈ 450, Pyr.
n.s.

**Supplementary Figure 2. More example LIP cells.**

Each column represents one example cell. Upper and middle panels, PSTH and choice divergence in the heading discrimination task. Same conventions as in Fig. 2. Cell depth (Dep.) was estimated from the spatial relationship between the gray matter pattern and the recording site (**Supplementary Figure 1**). Cell type (Pyr., pyramidal; Int., interneuron) was inferred from the bimodal distribution of spike waveform widths (data not shown). Lower panel, activities in the memory-guided saccade task. Solid and dashed curves represent average PSTHs when the saccade target was in and ouside the cell's response field, respectively. Inset, spatial tuning (mean ± s.e.m.) of the delay-period activity (25–900 ms from the target offset, gray shaded area); thick arrow, the preferred saccade direction derived from vector sum, with the number indicating its length (in Hz); ***, **, and n.s. indicate $p < 0.001$, $p < 0.01$, and $p > 0.05$, respectively, from one-way ANOVA.

**(a)** Two "typical" cells that had choice signals similar to the population average shown in Fig. 3.

**(b)** Two cells that were "non-typical" in the sense that their delay-period activities cannot predict their choice activities. Cell #7 had a strong delay-activity in the memory-guided saccade task (bottom panel) but exhibited little choice signal in the visual and combined heading discrimination task (top and middle panels), and vice versa for Cell #8.

**(c)** Three examples where choice signal of one modality (vestibular, visual, and combined, respectively) was more prominent than the other two. Colored circles are the same as in **Supplementary Figure 1**.

**(d)** One example cell that only had modality signal (divergence between blue and red curves in the top panel) but no choice signal (divergence between solid and dashed curves in the top panel).

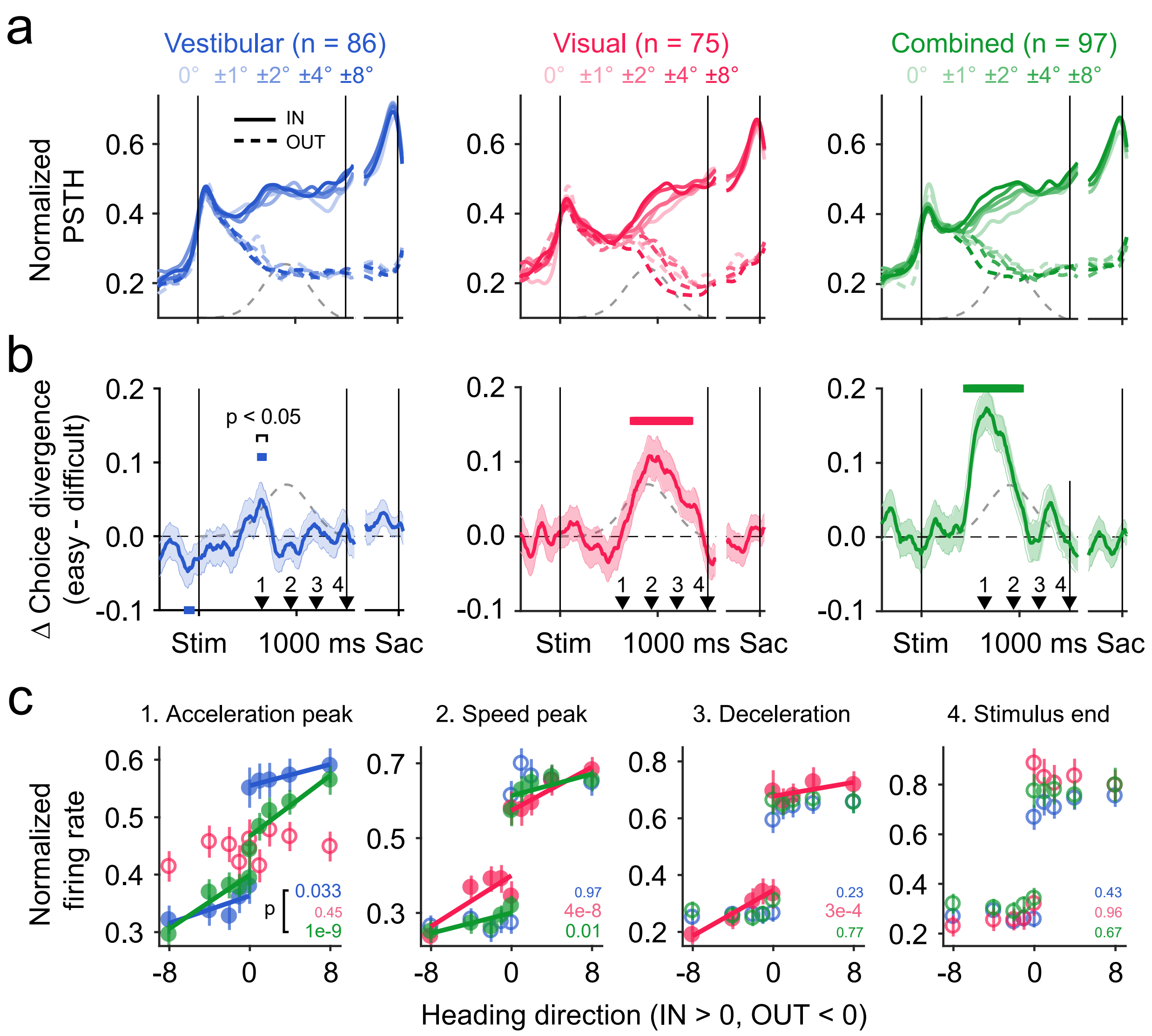


**Supplementary Figure 3. Task-difficulty dependence of choice signals**

**(a)** Population average of normalized PSTHs for different heading directions (the higher saturation, the larger headings). IN, trials toward the cell's response field (RF); OUT, trials away from the cell's RF. Only correct trials (but all trials for 0°) from cells with significant grand choice divergence (see **Methods**) under each condition were included in **a–c**.

**(b)** Difference between the average choice divergence of easy trials (4° and 8°) versus difficult trials (0°, 1°, and 2°). Error bands, s.e.m. Horizontal color bar, $p < 0.05$, two-tailed paired t-test.

**(c)** Population average of normalized activity is plotted against the heading angle at four time epochs (250-ms windows centered at four arrowheads in **b**). Tuning curves were flipped such that positive headings always correspond to directions toward the cell's RF. We fitted the response difference between IN and OUT choices as a linear function of the absolute value of heading (task-difficulty). The p-values of these linear fits are shown in each panel, with the filled data points denoting the cases where $p < 0.05$, namely, LIP response significantly depends on task-difficulty. The inclined lines were linear fits using data from IN and OUT choices separately. Error bars, s.e.m.

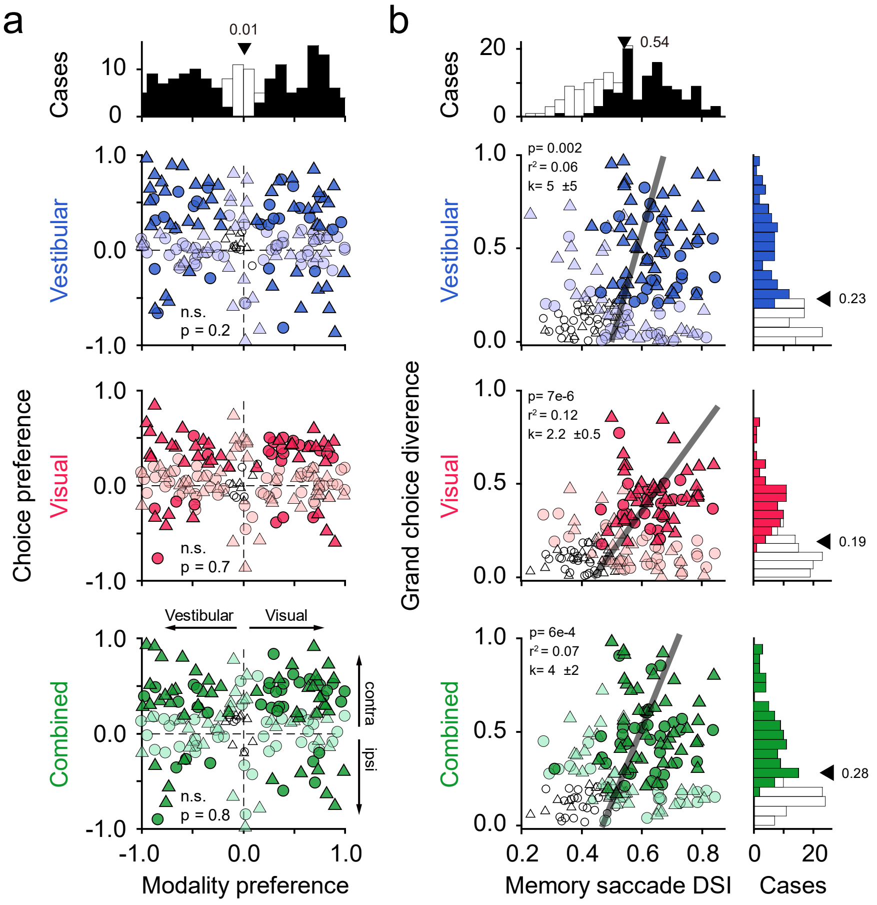


**Supplementary Figure 4. Macaque LIP is category-free**

**(a).** Choice signal was uncorrelated with modality signal. Ordinate, choice preference, which is similar to grand choice divergence (see **Methods**), except that its sign represents the location of the cell's RF (> 0, RF in the contralateral visual field; < 0, RF in the ipsilateral visual field). Abscissa, modality preference, which is similar to choice preference, except that the response difference between visual versus vestibular trials, instead of contra- versus ipsi-lateral choices, were compared. Circles, monkey P; triangles, monkey M. Shading indicates significance: dark color, both choice and modality preferences were significant ($p < 0.05$, two-sided permutation test, 1000 times); light color, either choice or modality preference was significant; open, none of them were significant. Top panel, histogram of modality preference; arrowhead, median. Note that for all three cue conditions, choice and modality preferences were uncorrelated (p-values from Pearson's correlation). This result is in keeping with a recent finding on rat PPC (**[11] Raposo, *et al.*, 2014)**.

**(b).** Choice signal was significantly, but only weakly, correlated with delay-period activity in the memory-guided saccade task. Ordinate, grand choice divergence, which is equal to the absolute value of choice preference in **a**. Abscissa, direction selective index (DSI) of spatial tuning in the memory-guided saccade task (see **Supplementary Figure 2**). DSI ranged from 0 to 1 and was defined as $\Delta R/(\Delta R + 2\sqrt{\text{SSE}/(N-M)})$, where $\Delta R$ is the difference between the maximum and minimum responses, SSE is the sum squared error, $N$ is the total number of trials, and $M$ is the number of target locations ($M = 8$). Conventions are the same as in **a**. Black thick lines and statistics were from type II regression. Note that for all three cue conditions, choice signal and memory-saccade activity were only weakly correlated (small $r^2$s). This result is in keeping with a recent finding on monkey LIP for visual decision making **([12] Meister, *et al.*, 2013)**.

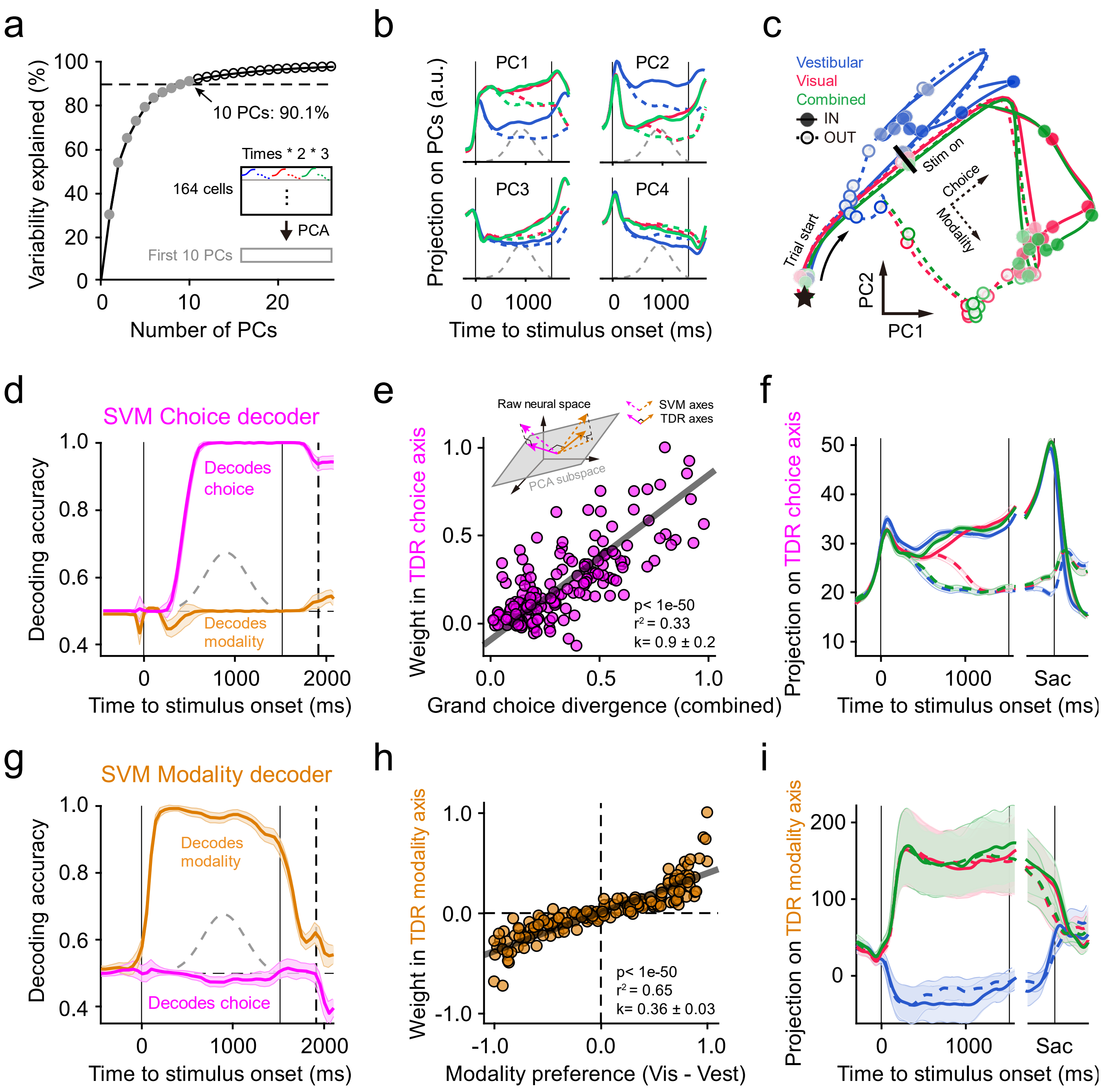


**Supplementary Figure 5. De-mixing of choice and modality signals**

**(a–c)** Principle component analysis (PCA). We first built an $n$ by $6t$ activity matrix ($n$, number of cells; $t$, number of time bins), in which each row was concatenated average responses grouped by all (six) possible combinations of cue condition and choice for each cell (**a**, inset, black rectangle). Then we reduced its dimensionality (number of rows) using PCA, resulting in a 10 by $6t$ matrix (**a**, inset, gray rectangle) with each row representing population activity projected on each principal component (PC).

**(a)** Percentage of variability explained is plotted against the number of PCs included. The first 10 PCs accounted for more than 90% variability of the original dataset.

**(b)** Population activities projected onto the first 4 PCs. Conventions are the same as before.

**(c)** Population trajectories in the plane spanned by PC1 and PC2. Black star, trial start; black bar, stimulus on. The time interval between two adjacent symbols on a same trajectory is 200 ms. Note that choice and modality signals evolved along two directions (dashed axes) that were nearly orthogonal but not aligned to the PC1-PC2 axes (solid).

**(d–i)** Targeted dimensionality reduction (TDR). To obtain de-noised and orthogonal axes of and task-relevant axes, we combined unsupervised PCA and a supervised method, linear support vector machine (SVM). Briefly, we trained two SVM decoders, a choice decoder (**d**) and a modality decoder (**g**), to decode the monkey's choice and the sensory modality, respectively, based on single-trial "pseudo-population" responses (**[11] Raposo*, et al.*, 2014)**. All spikes during the 1.5-s stimulus duration were used for training, and the resulting weights were averaged from 1000 bootstraps. Once trained, the decoders were tested for each 100-ms window (advancing in 50-ms steps) by another sets of pseudo-population responses (**d** and **g**). Finally, we projected the two SVM axes onto the PCA subspace, and then orthogonalized them by QR decomposition (**e**, inset) (**[13] Mante*, et al.*, 2013)**, yielding two de-noised and task-relevant axes, the TDR choice axis (**e** and **f**) and the TDR modality axis (**h** and **i**).

**(d)** Decoding accuracy of the SVM choice decoder when it was used to decode monkey's choice (purple) and stimulus modality (yellow), respectively. Error bands, standard deviation from 300 bootstraps of the test datasets.

**(e)** Each cell's weight in the TDR choice axis is plotted against its grand choice divergence. Gray line and statistics, type II regression. Inset, a schematic of the TDR process.

**(f)** Population activity projected on the TDR choice axis. Error bands, standard deviation from 1000 bootstraps of the whole TDR process.

**(g–i)** Same as in **(d–f)** but for the SVM modality decoder and the TDR modality axis.

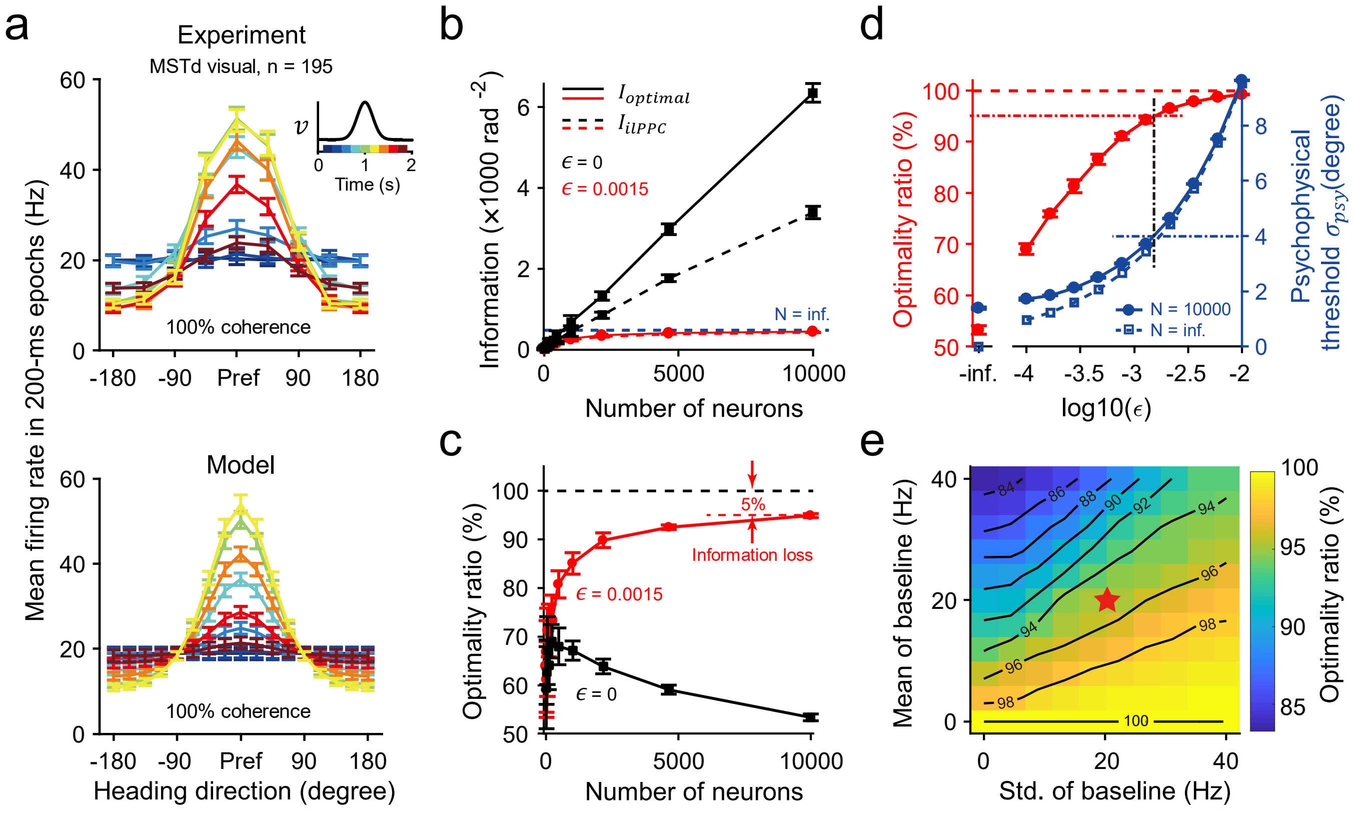


**Supplementary Figure 6. Information loss of ilPPC solution with heterogeneous MSTd population**

**(a).** Spatio-temporal tuning curves of real (upper panel) and modeled (lower panel) MSTd population. Tuning curves were derived from non-overlapping 200-ms time epochs (color coded; see inset) and aligned according to their preferred directions. Error bars, s.e.m.

**(b).** Fisher information as a function of the number of neurons $N$ that can be decoded from a heterogeneous MSTd population, with ($\epsilon = 0.0015$, red) or without ($\epsilon = 0$, black) differential correlations, by an optimal decoder (solid curves) or an ilPPC decoder (dashed curves). Blue dashed line indicates the saturated information when the number of neurons goes to infinity, computed with $\epsilon = 0.0015$ from Equation S11.

**(c).** Optimality ratio as a function of the number of neurons, $N$ under two differential correlation levels $\epsilon = 0.0015$ (red) and $\epsilon = 0$ (black). The optimality ratio is defined as the ratio of the information (solid curves in **b**) that can be recovered by the ilPPC solution (dashed curves in **b**) over the total information. For $N = 10000$ and $\epsilon = 0.0015$, the optimality ratio is around 95%, corresponding to around 5% information loss.

**(d).** The optimality ratio of ilPPC ($N = 10000$, red curve and left axis) and the psychophysical threshold $\sigma_{psy}$ (blue curve and right axis) as a function of $\epsilon$. The $\sigma_{psy}$ was calculated from Equations S6 and S15. Blue dashed curve is the saturated psychophysical threshold when $N$ goes to infinity, computed from Equations S11 and S15. Error bars in **b**–**d**, standard deviations from 10 runs.

**(e).** The optimality ratio of ilPPC as a function of the mean and the standard deviation of the baseline $B_i$. Note that the gradient around the biologically realistic regime (red star) is small, indicating that our estimated information loss of ~5% is reliable.

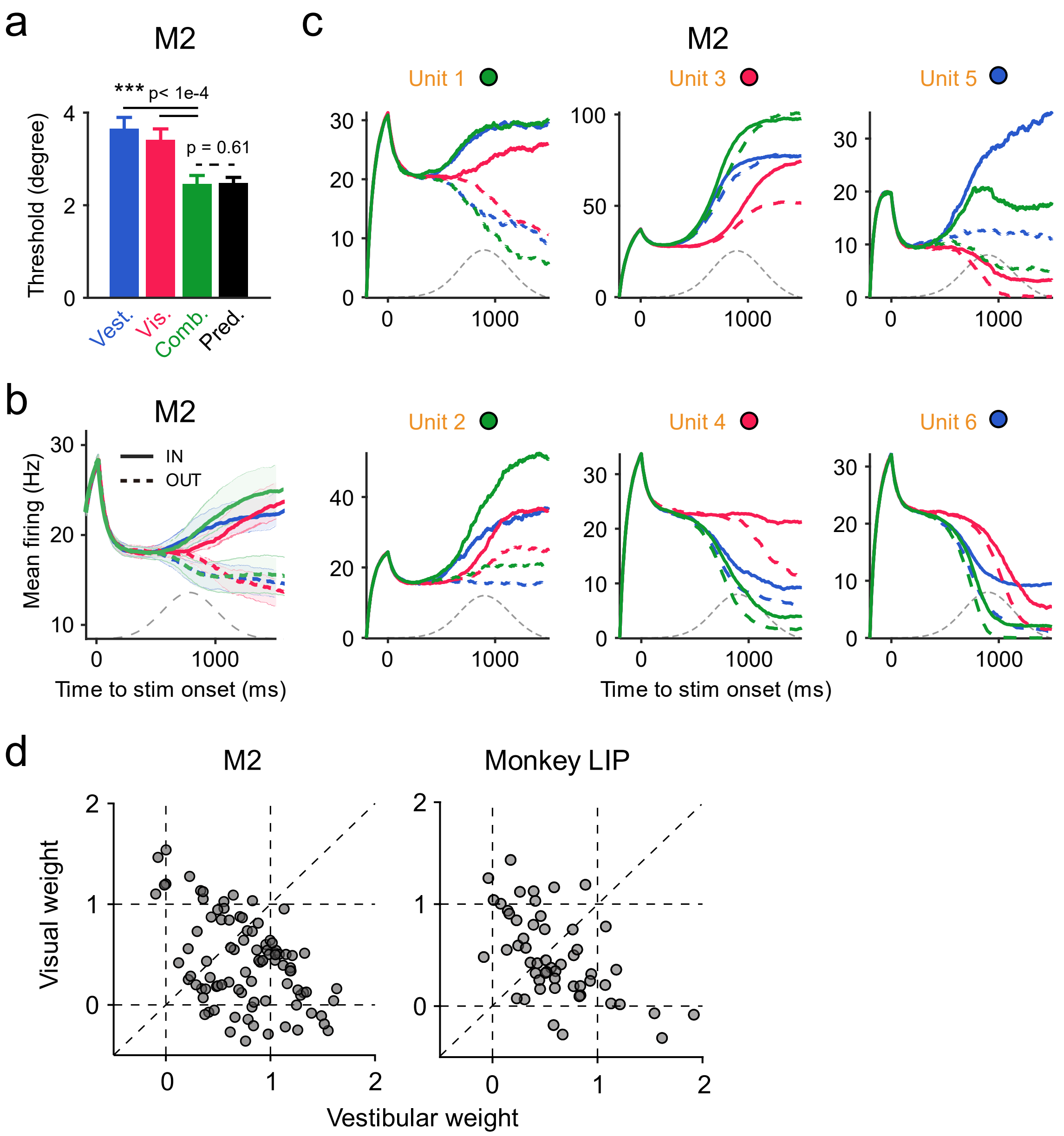


**Supplementary Figure 7. Model M2 achieves optimal behavior with heterogeneous units**

**(a)** M2 performed the task near-optimally, as did the monkeys and the homogeneous model M1.

**(b)** Mean firing rate of LIP units in M2 (n = 30). Error bands, s.e.m.

**(c)** Example LIP units showing heterogeneity in M2. For each unit, choice signal (the difference between solid and dashed curves) can be strongest in the combined condition (Unit 1 and 2), the visual condition (Unit 3 and 4), or the vestibular condition (Unit 5 and 6). Meanwhile, modality signal (the difference between blue and red curves, averaged across choices) can prefer either the visual cue (Unit 4 and 6) or the vestibular cue (Unit 3 and 5). Similar patterns have been found in our monkey LIP data (see **Figure 2** and **Supplementary Figure 2**).

**(d)** Visual and vestibular weights of units in Model M2 (left panel) and the LIP data (right panel) derived from the best fit of a linear weighted summation model $r_{combined} = w_{vestibular} r_{vestibular} + w_{visual} r_{visual} + C$. Each dot represents one neuron; only the neurons with significant fits are shown (linear model's p < 0.05).

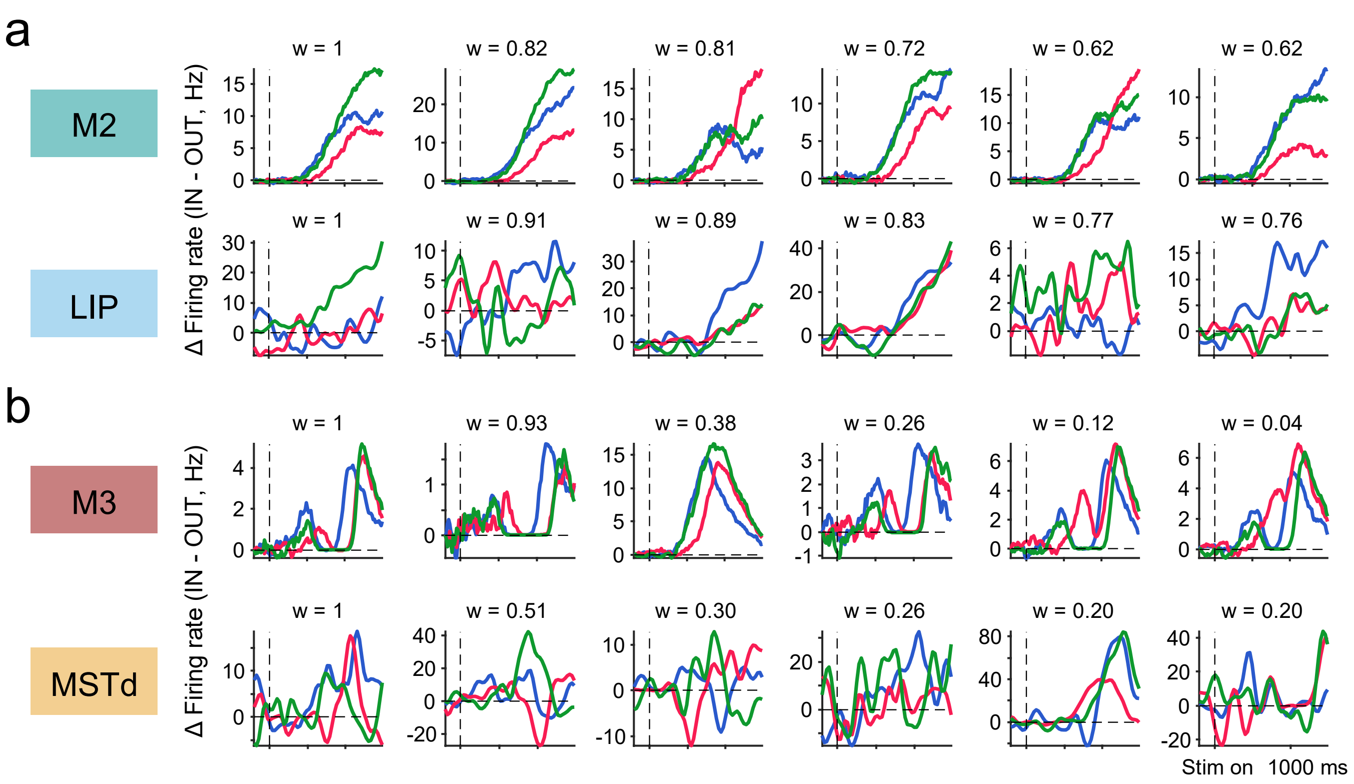


**Supplementary Figure 8. Example units in the linear reconstruction of M1**

**(a)** Six example units are shown ranked by their normalized readout weights in the linear fits (upper row, M2; lower row, LIP).

**(b)** Same as in (**a**) but for M3 (upper row) and MSTd (lower row).

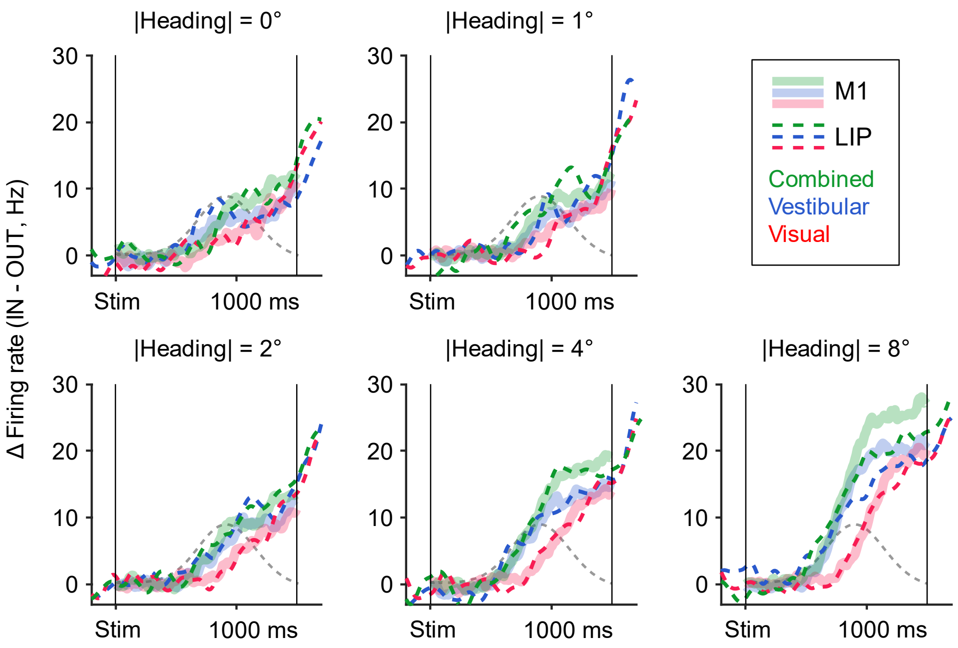


**Supplementary Figure 9. Linear reconstruction of M1 with cost function calculated from separated heading angles**
The same as **Figure 5d**, expect that the cost function used in the linear regression included error terms calculated from each heading angle separately (Equation 9 in the main text). The optimal traces from M1(thick shaded bands) and the linear readout of LIP (dashed curves) for each absolute value of heading angle (0°, 1°, 2°, 4°, 8°) are plotted in each panel. Note that although the cost function was calculated from each heading angle separately, each LIP cell still had only one readout weight in the linear regression (different headings shared the same $w_{\mathrm{LIP},i}$ in Equation 9).

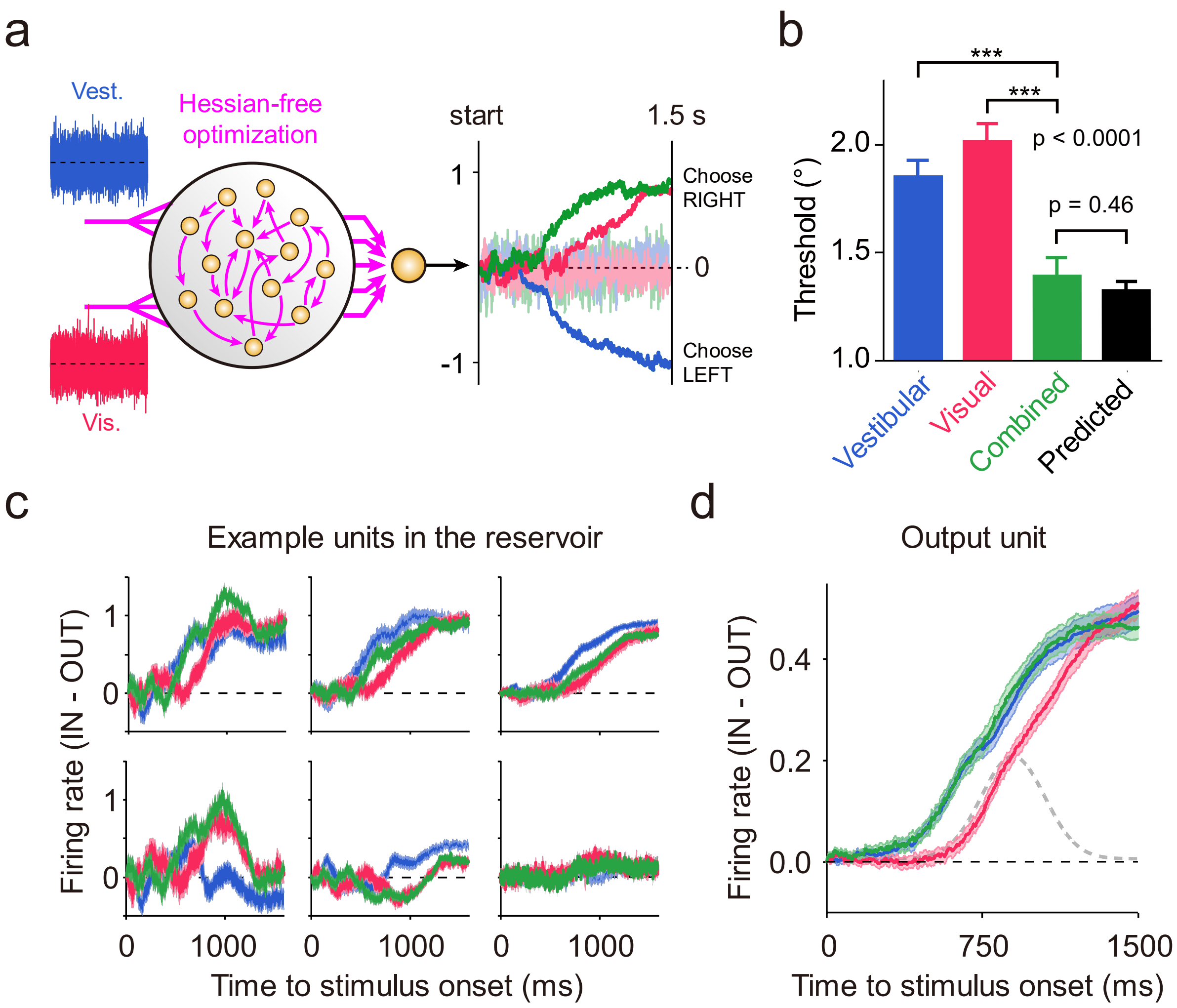


**Supplementary Figure 10. A trained recurrent neural network (RNN) performing multisensory decision-making task**

**(a)** The RNN receives noisy heading inputs from vestibular acceleration (top) and visual speed (bottom), processes the information by reverberating dynamics of the reservoir units (yellow dots inside the black circle), and uses the response of a linear readout unit (big yellow unit outside) to make heading judgment at the end of a trial (> 0, choose rightward; < 0, choose leftward). Before training, the synaptic weights of the RNN were randomly initiated, giving rise to noisy responses of the output unit (light-colored traces fluctuating around the dashed line) and chance-level behavioral performance. Then we trained the RNN with Hessian-free optimization (**[14] Martens, 2010; [15] Martens and Sutskever, 2011; [13] Mante*, et al.*, 2013)**. By doing this, we optimized all the synaptic weights in the RNN (purple arrows) to minimize the sum squared error between the output traces and the "ideal traces" defined as the time integrals of the corresponding noiseless input signals. After training, the network could effectively suppress the noise and integrate the sensory inputs across time and cues (dark-colored traces).

**(b)** The trained RNN combined cues near-optimally. Conventions are the same as before. P-values, t-test, 25 repetitions.

**(c)** Responses of six example units in the reservoir. Trials were from 8° headings under three cue conditions; error band, s.e.m. Note that due to the inherent randomness of RNN, these units show a wide range of response patterns.

**(d)** Responses of the output unit. Error band, s.e.m.; gray bell-shape curve, the same speed profile used in the experiments.
